# Absence of postsynaptic activity on developing neurons alters gene expression profiles without preventing the refinement of inputs

**DOI:** 10.1101/2020.10.21.349142

**Authors:** Yumaine Chong, Ellis Cooper

## Abstract

Synaptic activity plays several roles as developing neurons make connections with their targets. It acts locally at synapses to influence the expression of genes needed to establish and maintain synaptic contacts. And, downstream it provides the necessary activity to strengthen and refine connections. Many studies have demonstrated how synaptic activity alters synaptic strength and increases synapse numbers. Much less is known, however, about the long-term consequences when a circuit develops without synaptic activity. To address this, we developed a mosaic model of sympathetic ganglia where synaptically-active and synaptically-inactive sympathetic neurons develop side-by-side *in vivo.* This model allowed us to address two issues. One is the relationship between activity and the refinement of converging inputs; the second is how synaptic activity contributes to a neuron’s gene expression profile. Our results indicate that converging presynaptic inputs to synaptically-silent neurons do not require postsynaptic activity to refine, provided these neurons share targets with synaptically-active neurons. Second, we demonstrate with single-cell RNA sequencing experiments that the expression of many genes by sympathetic neurons is independent of endogenous activity or local signals immediately downstream of excitatory postsynaptic potentials. An exception are genes required for neurotransmitter metabolism: We found that for a large sub-population of sympathetic neurons, synaptic activity increases the expression of adrenergic genes and supresses the expression of cholinergic genes. We conclude that signals generated locally at synapses do not initiate refinement of converging inputs, and that synaptic activity’s influence on a neuron’s gene expression profiles is complex and depends on context.

## Introduction

As neural circuits begin to form, the resulting synaptic activity plays a crucial role in shaping the patterns of connections while these circuits mature. A good example is the refinement of converging inputs, a process that occurs widely throughout the developing nervous system. The prevailing view is that the refinement of inputs during development depends critically on synaptic activity. The general consensus is that refinement is a competitive process: Converging inputs compete for molecules secreted in an activity-dependent manner from postsynaptic neurons; the successful inputs persist and are strengthened, and less successful ones are eliminated (Changeux and Danchin, 1976; Purves and Lichtman, 1980; Katz and Shatz, 1996; Lichtman and Colman, 2000). This view is supported by several studies that have blocked synaptic transmission, either pharmacologically or genetically, at specific synapses and examined the refinement of inputs on postsynaptic neurons (Thompson et al., 1979, Balice-Gordon and Lichtman, 1994; Buffelli et al., 2003; Kano and Hashimoto, 2009). Using such approaches, several studies indicate that inputs do not refine without activity and provide insight into molecular mechanisms on how postsynaptic neurons respond, in general, to synaptic activity (Buffelli et al., 2003; Cohen-Cory, 2002; Flavell and Greenberg, 2008; Hua and Smith, 2004; Redmond, 2008; Thompson et al., 1979; Wong and Ghosh, 2002).

Consistent with the above model for refinement of convergent inputs, we showed recently, using sympathetic ganglia with a deletion in the alpha 3 nicotinic receptor subunit gene (α3 KO), that developing preganglionic inputs to sympathetic neurons fail to refine when there was no synaptic transmission between preganglionic axons and postsynaptic sympathetic neurons (Chong et al., 2018). These results are in line with postsynaptic activity being necessary for inputs to refine. In contrast, when we enhanced mRNA translation by deleting the mRNA elongation factor binding protein, 4E-BP in α3 KO mice (double α3/4E-BP KO), the converging preganglionic inputs refined without postsynaptic activity (Chong et al., 2018). These latter results conflict with the widely-held belief that refinement of converging inputs depends critically on synaptic activity. Instead, they raise the possibility that refinement is not initiated by postsynaptic activity, but that non-cell autonomous mechanisms may be involved.

It was difficult for us to determine which cells initiated the refinement in double α3/4E-BP KO ganglia because the 4E-BP1/2 genes were knocked out globally, and therefore, in addition to postsynaptic neurons, several different cell types with enhanced mRNA translation could have initiated inputs to refine. Another consideration, blocking of synaptic transmission in autonomic ganglia disrupts activity of autonomic reflexes, and such disruption during postnatal development may influence how postganglionic neurons are innervated by preganglionic axons. We reasoned that a more definitive test of whether refinement of converging inputs depends critically on synaptic activity during development would be to compare neurons that had developed without synaptic transmission to neurons in the same sympathetic ganglion that were synaptically active, thereby keeping autonomic reflexes largely intact. If synaptic activity is necessary for converging inputs to refine, then one would expect that the preganglionic axons innervating synaptically-active neurons would refine as normal, but those axons that innervate neurons without synaptic transmission would not.

To test this predication, and by inference, that postsynaptic activity initiates refinement of converging inputs, we genetically engineered mice to have mosaic autonomic ganglia where half the neurons had a deletion in the α3 gene to eliminate all functional postsynaptic nAChRs and consequently developed without synaptic transmission, and the other half, which were randomly intermingled with the first, had functional postsynaptic nAChRs and were synaptically excited by preganglionic neurons. Mosaic sympathetic ganglia in these mice enabled us to compare directly the development and innervation of neurons that received endogenous synaptic activity over the first 1-2 postnatal months to neighbouring neurons that did not receive excitatory synaptic transmission and remained synaptically silent.

We used our model to address two issues. One is whether refinement of converging inputs can occur without postsynaptic activity. The second relates to a general issue of gene expression profiles on developing neurons. Gene expression profiles are often used to define the identity of different neuronal subtypes; yet, each neuron’s profile underlies both its cell-type identity and its cellular activities, making it difficult to determine the contribution of each. Using mosaic ganglia together with single-cell RNA sequencing (scRNAseq), we determined how synaptic activity contributes to a neuron’s gene expression profile by comparing profiles of neurons that receive endogenous synaptic activity to profiles of neurons that remained synaptically silent.

In contrast to the prevailing view, we found that preganglionic axons refined on both synaptically-active and synaptically-silent neurons, indicating that postsynaptic activity does not initiate the refinement of converging inputs. Moreover, our results point to a role for target activity in the refinement of upstream synapses. With respect to gene-expression profiles, we found many genes whose expression changed little between synaptically-active and synaptically-silent neurons, indicating that the expression of these genes is governed by non-activity-dependent mechanisms. On the other hand, we found that many synaptically-silent neurons in mosaic ganglia had large increases in the expression of cholinergic genes, indicating that to maintain an adrenergic phenotype, sympathetic neurons require synaptic activity.

## Results

Our experiments involve mice with mosaic sympathetic ganglia composed of synaptically-silent neurons that developed side-by-side with synaptically-active neurons. We accomplished this by exploiting random X-chromosome inactivation (see Materials and Methods). Briefly, we used the human ubiquitin promoter to express from the X chromosome either the nAChR subunit gene α3 (referred to as X^α3^), or the gene for mRFP1 (referred to as X^RFP^). We then mated these mice to α3 KO mice (mice with a deletion in the endogenous α3 gene on chromosome 9 (Krishnaswamy and Cooper, 2009; Rassadi et al., 2005; Xu et al., 1999)). The phenotype of these progeny was phenotypically normal, yet, due to random X-inactivation, females had mosaic sympathetic ganglia in which only neurons containing X^α3^ expressed functional nAChRs, whereas neurons containing X^RFP^ had no nAChRs (**Figures 1A and B; Figure S1**).

**Figure 1.**
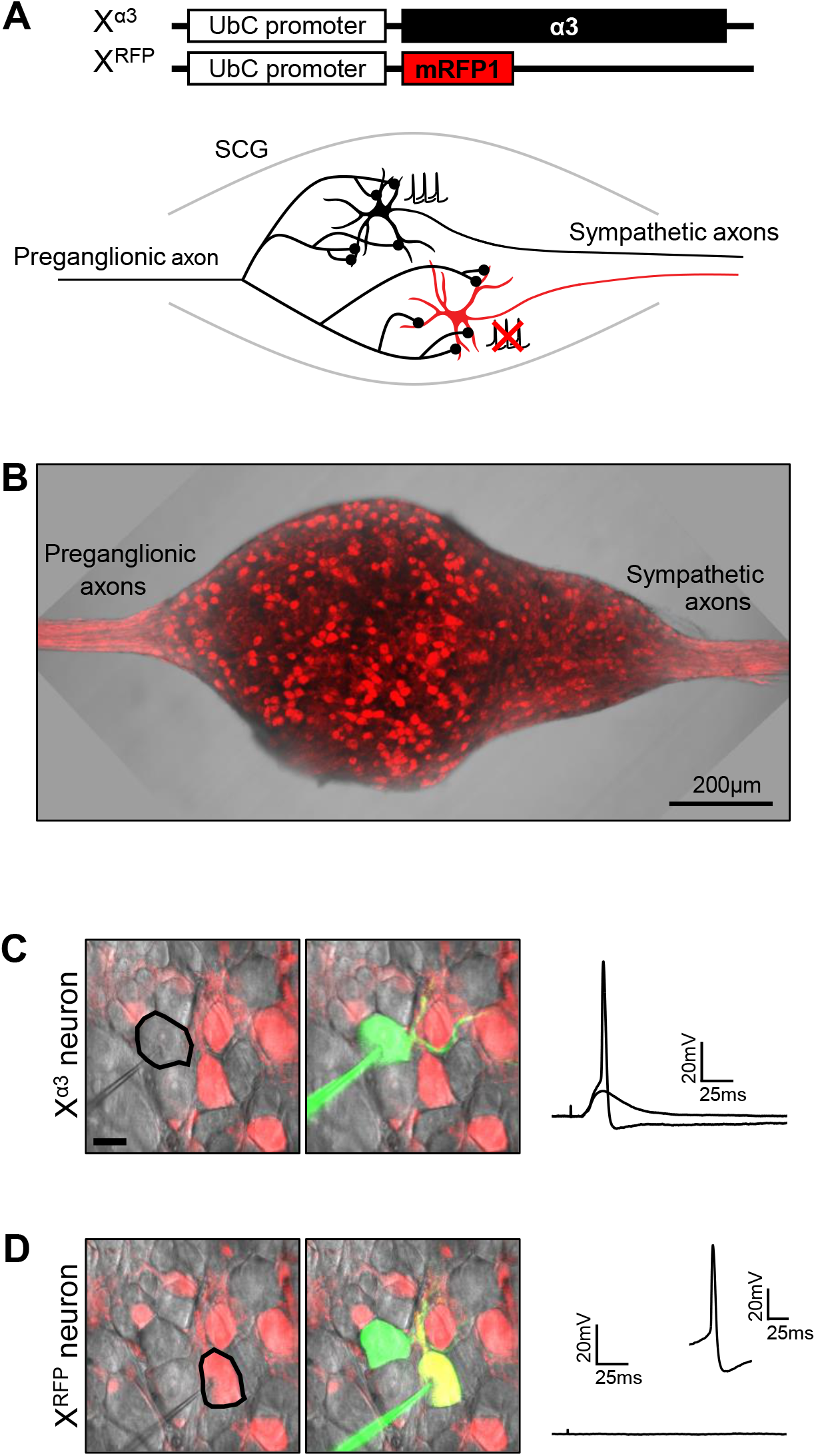
Random X inactivation generates mosaic population of synaptically-active and inactive neurons in sympathetic ganglia. (A) Schematic drawing of a mosaic SCG with a preganglionic axon innervating a neuron (black) with α3-containing nAChRs (Xα3) and a neuron (red) without nAChRs (XRFP) that expresses RFP. See also Figure S1. (B) Superimposed DIC and confocal images of an intact mosaic SCG composed of non-fluorescent Xα3 neurons and RFP-expressing XRFP neurons. See also Figure S1. (C, D) Superimposed DIC and confocal images (scale bar, 2Oμm) of an Xα3 (C) and an XRFP (D) recorded intracellularly with a sharp electrode filled with Alexa Fluor 488 hydrazide (green) and the corresponding electrophysiological response to preganglionic nerve stimulation (right). Preganglionic stimulation evokes suprathreshold EPSPs from Xα3 neurons and no responses from XRFP neurons. Inset in D shows an action potential from a XRFP neuron evoked with current injection.

To demonstrate that sympathetic ganglia in these mice were indeed mosaic, we recorded from hundreds of neurons in intact superior cervical ganglia (SCG). Only X^α3^ neurons were excited by preganglionic nerve stimulation (**Figure 1C**). Moreover, the fast EPSPs on these X^α3^ neurons were indistinguishable in amplitude and time course from those on age-matched sympathetic neurons from wild-type mice. On the other hand, stimulating the preganglionic nerve failed to evoke any detectable responses from X^RFP^ neurons, although these X^RFP^ neurons received morphological contacts (**Figure S1**) and were capable of generating overshooting action potentials with direct current injection (**Figure 1D**), as was reported previously for sympathetic neurons in α3 KO mice (Krishnaswamy and Cooper, 2009; Chong et al., 2018).

Before using these mosaic ganglia to investigate whether refinement of converging inputs can occur without postsynaptic activity, or how the absence of synaptic activity influences gene expression profiles, we characterized the morphology of synaptically-active X^α3^ neurons and synaptically-silent X^RFP^ neurons, and the innervation of these neurons by preganglionic axons. Since X^α3^ neurons receive functional innervation from preganglionic axons, we expected that the dendrites on X^α3^ neurons would be similar to those on SCG neurons in wild-type mice. And indeed, we show that synaptically-active X^α3^ neurons had elaborate dendritic arbors: the average total dendritic outgrowth (TDO) and the number of primary dendrites were not statistically different from P28 SCG neurons in wild-type mice (**Figures 2A, C and D**).

**Figure 2.**
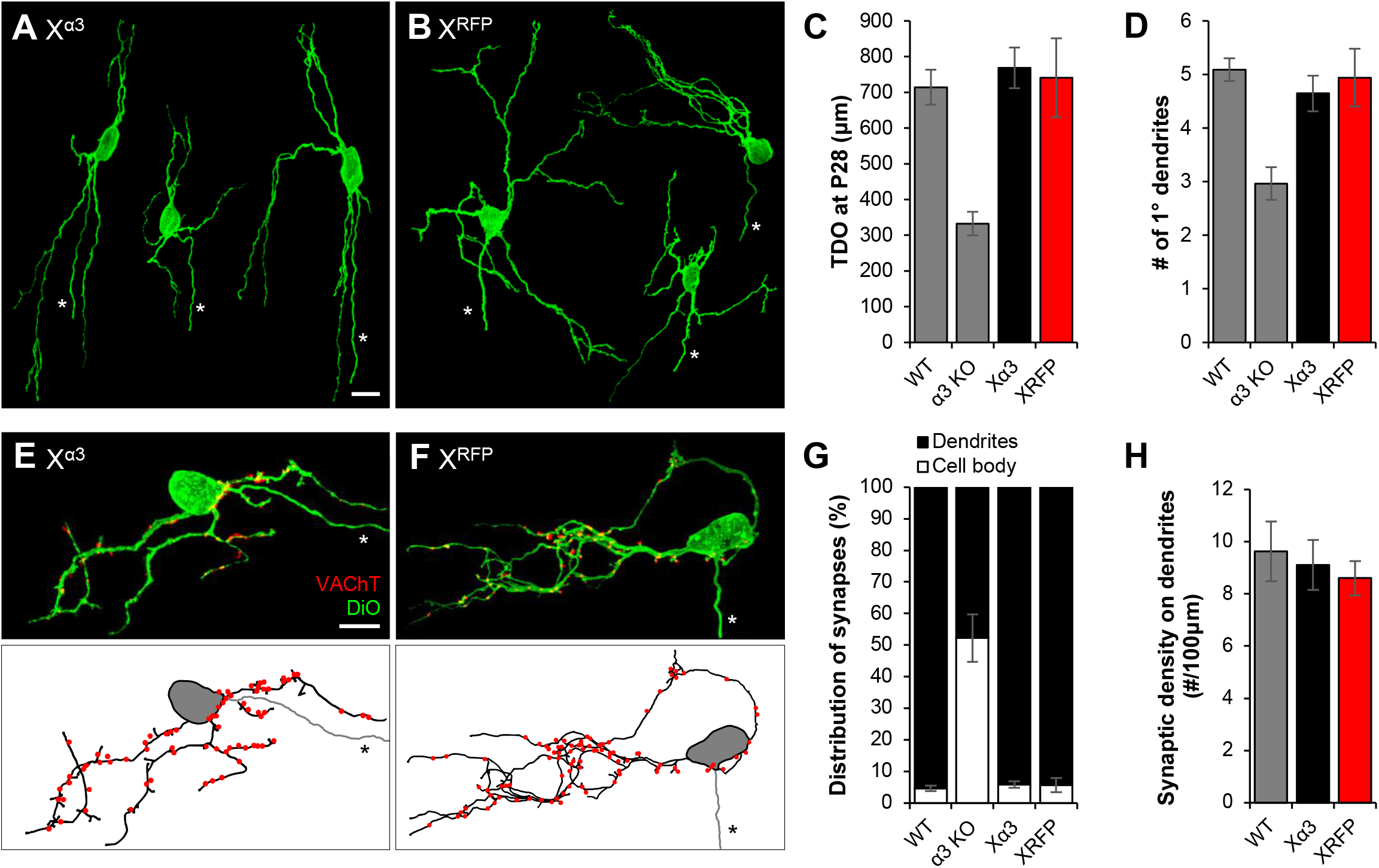
Dendritic growth and synaptic targeting on Xα3 and XRFP neurons in mosaic SCG. (A-B) Maximum intensity projections of DiO-labelled (A) Xα3 neurons and (B) XRFP neurons from mosaic SCG labelled at P28. All neurons are shown at the same scale; axons are marked by an asterisk. Scale bar, 2Oμm. In each panel, neurons are from different ganglia and have been tiled for comparison. (C-D) Quantification of (C) average total dendritic outgrowth (TDO) per neuron at P28, and (D) average number of primary dendrites per neuron at P28. Black columns represent Xα3 neurons and red columns represent XRFP neurons. Grey columns indicate average values from P28 WT and α3 KO neurons for comparison. WT and α3 KO data are from Chong et al., 2018. (E-F) Top: Maximum intensity z-projection of a DiO-labelled (green) (E) Xα3 neuron and (F) XRFP neuron in P28 mosaic SCG, immunostained for VAChT (red); axons are marked by an asterisk. VAChT puncta not touching the neuron were removed for clarity. Scale bar, 2Oμm. Bottom: Skeletonized reconstructions showing dendritic arbors (black), axon (grey, marked by an asterisk), and preganglionic axon varicosities (red) determined by VAChT staining. (G) Average distribution of varicosities on the cell body (open) and dendrites (filled) of Xα3 neurons and XRFP neurons in mosaic SCG at P28. WT and α3 KO data are from Chong et al., 2018. (H) Average number of synapses per 100 μm length of dendrites on XRFP, Xα3 and wild-type SCG neurons at P28. For C, D, G, H error bars represent ± SEM. For Xα3, n=10 neurons, and for XRFP, n=10 neurons (8 mice).

Previously, using SCG from α3 KO mice, we demonstrated that sympathetic neurons that develop without synaptic transmission had stunted dendrites (Chong et al., 2018). Therefore, we were expecting that dendritic growth on X^RFP^ neurons would be significantly less than that on X^α3^ neurons. Surprisingly, synaptically-silent X^RFP^ neurons in mosaic SCG had dendritic outgrowth that closely matched that of active X^α3^ neurons and wild-type neurons, both in total dendritic outgrowth (TDO), as well as the number of primary dendrites, and significantly greater than SCG neurons in α3 KO mice (**Figures 2B, C and D**).

Our previous work showed that the phosphorylation of the mRNA translation repressor, 4E-BP is significantly reduced in SCG neurons in α3 KO mice compared to age-matched controls (Chong et al., 2018). Moreover, in mice where 4E-BP had been genetically removing 4E-BP from α3 KO mice (4E-BP/ α3 DKO), dendritic outgrowth on SCG neurons is not significantly different from wild-type SCG neurons. Relevantly, in mosaic SCG, we show that phosphorylated-4E-BP levels in X^α3^ were not statistically different from X^RFP^ neurons (**Figure S1**), consistent with cap-dependent translation overriding the effects of synaptic inactivity on dendritic growth.

To determine the innervation of X^α3^ and X^RFP^ neurons by preganglionic axons, we labelled P28 neurons in mosaic SCG with DiO and stained for the vesicular acetylcholine transporter (VAChT), a presynaptic marker localized at synapses. Over 90% of synapses on X^α3^ neurons were targeted to dendrites, not statistically different from those on SCG neurons in wild-type mice (**Figures 2E, F and G**).

We showed previously that in the absence of synaptic transmission preganglionic axons establish and maintain morphological synapses on SCG neurons in α3 KO mice that appear normal (Krishnaswamy and Cooper, 2009); however, more than 50% of the presynaptic terminals targeted silent synapses to the soma of SCG neurons (Chong et al., 2018). In contrast, we found that over 90% of the silent synapses on X^RFP^ neurons in mosaic SCG were targeted to the dendrites, a proportion similar to those on X^α3^ neurons, and on SCG neurons in wild-type mice (**Figures 2E, F and G**). The total number of synapses on SCG neurons increases ~2-fold over the first postnatal month (Krishnaswamy and Cooper, 2009; Lehigh, West and Ginty, 2017), and we found that there is no statistical difference in synapses per length of dendrites between X^RFP^, X^α3^ and wild-type SCG neurons at 1 month (**Figures 2H**).

### Refinement of converging preganglionic axons in the absence of postsynaptic activity

Having established that that X^α3^ and X^RFP^ neurons in mosaic ganglia had normal dendritic morphologies and were innervated by preganglionic axons, we asked whether converging preganglionic axons contacting SCG neurons undergo refinement in the absence of synaptic transmission. To quantify refinement, we recorded jumps in the evoked EPSPs on SCG neurons while gradually increasing the stimulus to the preganglionic nerve (Chong et al., 2018). A few days after birth, X^α3^ neurons were functionally innervated by 6-8 preganglionic inputs. However, by one month, most inputs were eliminated and only 1-3 remained (**Figures 3A and B**), the same as the refinement of preganglionic axons innervating SCG neurons in wild-type mice (**Figure 3E**) (Chong et al., 2018).

**Figure 3.**
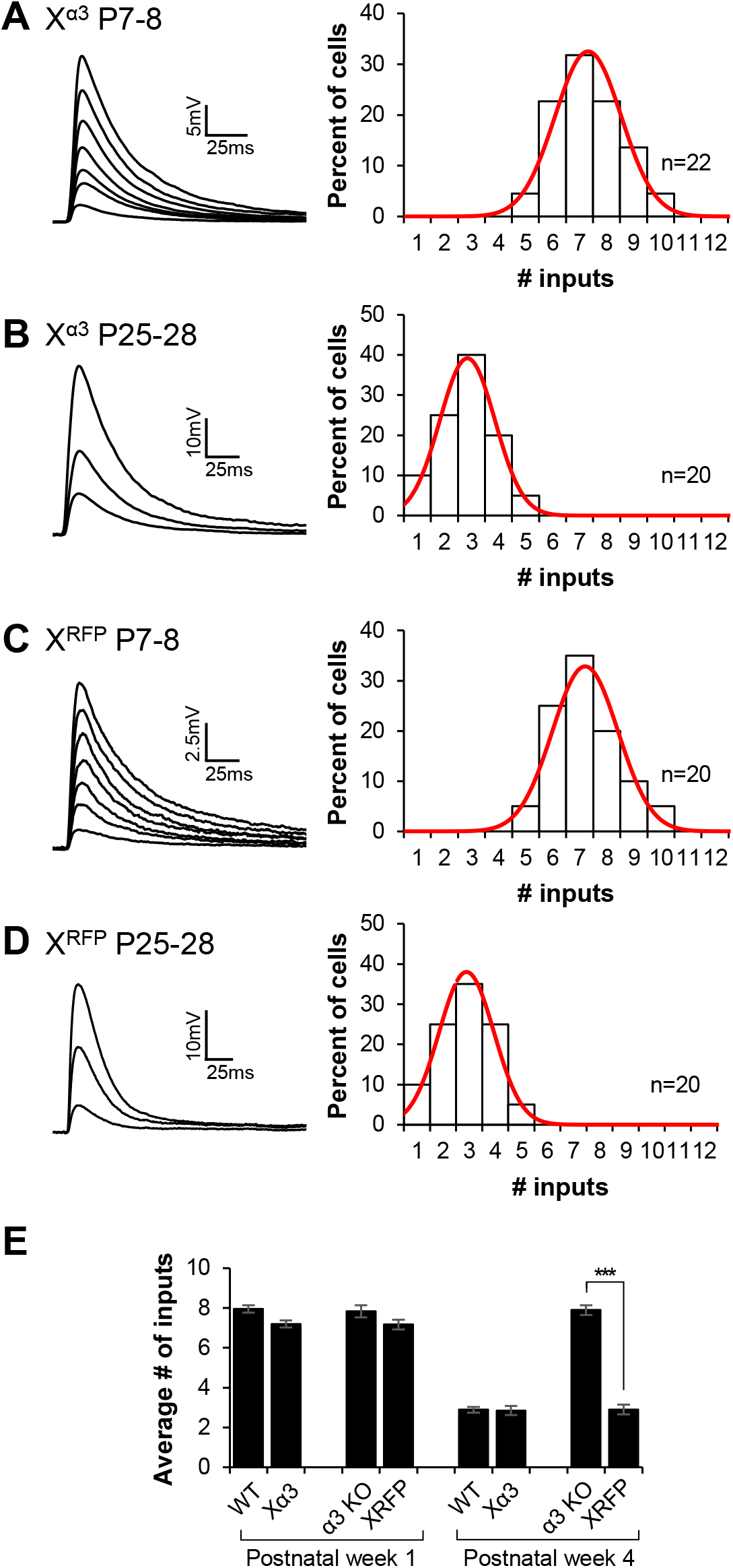
Refinement of converging preganglionic axons on synaptically-silent neurons in mosaic SCG. (A-D) Left: Representative EPSPs in a (A) P7-8 Xα3 neuron, (B) P25-28 Xα3 neuron, (C) P7-8 XRFP neuron, and (D) P25-28 XRFP neuron in mosaic SCG evoked by increasing stimuli to the preganglionic nerve. Right: Distribution of SCG neurons innervated by the number of inputs, fit with a Gaussian function. Each distribution in A-D contains data from at least 8 mice; n refers to the number of neurons. (E) The average number of axons innervating WT neurons, α3 KO neurons, Xα3 neuron, and XRFP neurons at postnatal week 1 and 4. WT and α3 KO data are from Chong et al., 2018. Error bars represent ± SEM; ***p<0.001

To quantify refinement on X^RFP^ neurons, 2 day prior to recording, we unsilenced synapses by infecting mice with adenoviruses containing the α3 nAChR subunit gene (Krishnaswamy and Cooper, 2009; Chong et al., 2018). X^RFP^ neurons were similarly innervated by 6-8 preganglionic axons a few days after birth (**Figure 3C**). Surprisingly, however, by one month, X^RFP^ neurons were innervated by 1-3 preganglionic axons, the same number innervating X^α3^ or wild-type neurons at P28 (**Figures 3D and E**). These data demonstrate that converging preganglionic axons do not require postsynaptic activity to refine.

The refinement on X^RFP^ neurons was unexpected in view of our previous work with SCG in α3 KO mice (Chong et al., 2018). Synaptic transmission is completely abolished on both X^RFP^ neurons and SCG neurons in α3 KO mice (Chong et al., 2018; Krishnaswamy and Cooper, 2009), yet preganglionic axons converging on X^RFP^ neurons refine, whereas those innervating SCG in α3 KO mice do not (**Figure 3E**). A possible explanation for this difference in refinement between preganglionic axons in mosaic SCG and those in SCG of α3 KO mice could be a difference in the innervation of targets that mediate sympathetic reflexes. In α3 KO mice, for example, there is no synaptic transmission in sympathetic ganglia and sympathetic reflexes are substantially reduced (Xu et al., 1999; Campanucci, Krishnaswamy and Cooper, 2010). On the other hand, in mosaic SCG, the targets are functionally innervated by X^α3^ neurons and autonomic reflexes appear normal. Therefore, we asked whether there were differences in target innervation between synaptically-silent X^RFP^ neurons and SCG neurons α3 KO mice.

### Sympathetic neurons in mosaic mice and α3 KO mice innervate the iris differently

We focused on the radial muscles of the iris, a tissue that causes pupil dilation upon sympathetic nerve stimulation (Hill et al., 1991). Pupillary reflexes are severely disrupted in α3 KO mice (Xu et al., 1999), whereas in mosaic mice, the reflex appears normal.

First, we determined the number of SCG neurons projecting to the iris, and then we quantified the density of sympathetic innervation. To determine the number of SCG neurons, one-month-old mice were injected with the retrograde tracer, cholera toxin subunit B conjugated to Alexa Fluor 488 (CTB-488) into the anterior chamber of one eye to back-label neurons in the SCG. We found that 35 ± 3.56 (mean ± s.e.m) SCG neurons projected to the iris in wild-type mice at P28, less than 0.5% of the total number of neurons in the ganglion (**Figures 4A and D**). These neurons were mostly distributed in the rostral end of the SCG and sent their axons exclusively to the ipsilateral eye, similar to iris innervation in the rat (Luebke and Wright, 1992). In age-matched α3 KO mice, the number of SCG neurons projecting to the iris was not statistically different from that in wild-type mice (37 ± 1.11; **Figures 4B and D**), and as in wild-type mice, most were localized to the rostral end and innervated the iris on the ipsilateral side. Moreover, in mosaic ganglia, the total number of neurons projecting to the iris was not statistically different from that in wild-type or α3 KO mice (35 ± 4.23; **Figures 4C and D**); relevantly, about half of these were X^RFP^ neurons (**Figure 4E**). These data indicate that innervation of iris muscle does not depend on whether neurons are synaptically active or synaptically silent.

**Figure 4.**
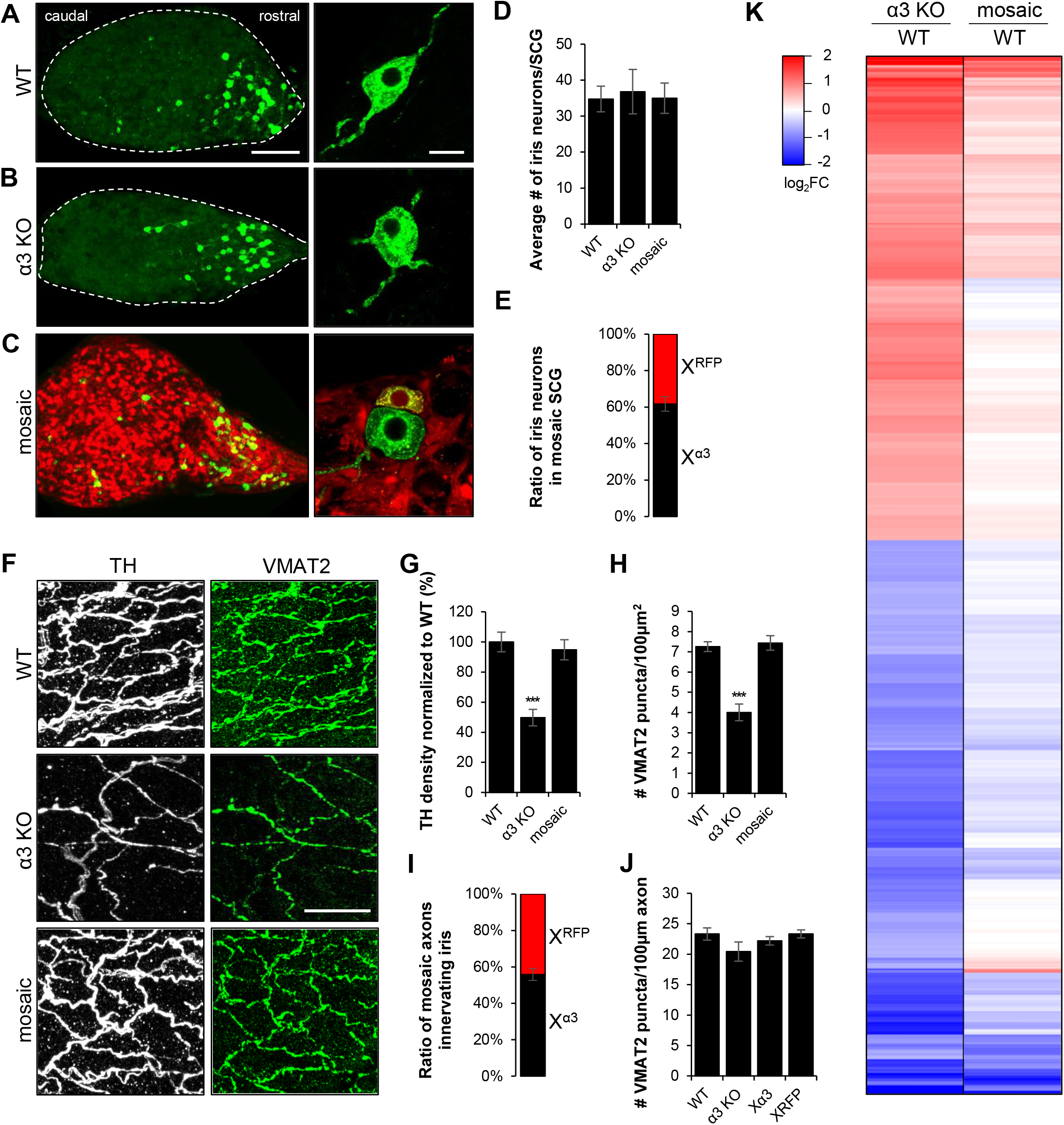
Innervation density of the iris is lower in α3 KO mice than in WT mice, yet X^α3^ neurons and X^RFP^ neurons in mosaic SCG innervate the iris at roughly similar proportions. (**A-C**) Left: Maximum intensity projection of CTB-488-labelled neurons (green) in intact SCG. Neurons were retrogradely labelled from the iris in (**A**) WT, (**B**) α3 KO, and (**C**) mosaic mice at P28. Scale bar, 200μm. Right: CTB-488-labelled neurons (green) at higher magnification. Scale bar, 20μm. (**D**) Average number of neurons innervating the iris per WT, α3 KO and mosaic SCG at P28. Error bars represent ± SEM. For WT, n=8 SCG (8 mice), for α3 KO, n=7 SCG (7 mice), and for mosaic, n=9 SCG (9 mice). (**E**) Average ratio of CTB-488-labelled X^α3^ neurons (black) and X^RFP^ neurons (red) per SCG innervating the iris in mosaic mice at P28, n=9 SCG (9 mice). (**F**) Maximum intensity projections of immunostaining for TH (white; left) and VMAT2 (green; right) in iris from WT, α3 KO, and mosaic mice at P28. Scale bar, 20 μm. (**G**) Average area of the iris innervated by TH-positive axons in WT, α3 KO and mosaic mice at P28, normalized to WT. (**H**) Average number of VMAT2 puncta per 100μm2 square area of iris in WT, α3 KO and mosaic mice at P28. (**I**) Average ratio of TH-positive axons from X^α3^ neurons (black) and X^RFP^ neurons (red) per iris in mosaic mice at P28. (**J**) Average number of VMAT2 puncta per 100μm length of skeletonized TH-positive axons from WT, α3 KO, X^α3^ or X^RFP^ SCG neurons innervating the iris at P28. (**K**) Heatmap generated from bulk RNA sequencing showing a random subset (800 of 6000) of differentially expressed genes (Log2FC>1.5, p<0.05) between WT and α3 KO irises (left), and between WT and mosaic irises (right). See also Table 1. For D, E, G, H, I and J, error bars represent ± SEM; ***p<0.001. For G-J, WT, n=10 irises (5 mice), and for α3 KO, n=10 irises (5 mice), and for mosaic, n=8 irises (4 mice).

Next, we asked whether the density of innervation of the iris by sympathetic axons in mosaic SCG differed from those in α3 KO mice. To quantify iris innervation, we immunostained iris muscle in wholemount for tyrosine hydroxylase (TH), a marker for sympathetic axons, and the vesicular monoamine transporter 2 (VMAT2), a marker for noradrenergic presynaptic varicosities. The axons from sympathetic neurons in mosaic SCG branched extensively to form a network of axons over the iris. The total density of axons was not statistically different from wild-type control (**Figures 4F, G and H**), and the density of X^RFP^ axons was approximately equal to that from X^α3^ axons (**Figure 4I**). On the other hand, the density of axons from α3 KO SCG neurons was approximately 50% of mosaic SCG or wild-type (**Figures 4F, G and H**). These results suggest that functional innervation of the target by sympathetic axons plays an important role in determining target innervation density.

We also quantified axonal presynaptic varicosities to the iris by immunostaining for VMAT2. When normalized to axon length, we found no statistical difference in the number of VMAT2 puncta along wild-type, α3 KO, X^α3^ and X^RFP^ axons innervating the iris (**Figure 4J**). This indicates that the appearance of noradrenergic presynaptic varicosities along sympathetic axons innervating targets is largely independent of action potentials conduction from the soma to the terminals. Likely, the induction of presynaptic varicosities relies on local axon-target interactions.

### Difference in expression of iris muscle genes between mosaic and α3 KO mice

In addition to differences in the density of axons innervating the iris, we investigated the activity of the targets. We expect that the targets of mosaic SCG neurons are active because they are functionally innervated by X^α3^ neurons, whereas targets activity in α3 KO mice is substantially reduced or absent.

To test this, we examined differences in iris muscle activity in mosaic mice and α3 KO mice. Long-term muscle activity is known to influence the expression of several muscle genes (Pillon et al., 2020; Kostrominova et al., 2005; Tang et al., 2000). Therefore, we used differences in gene expression as an effective readout of cumulative iris muscle inactivity over the first postnatal month *in vivo.* If activity of the iris in α3 KO mice is reduced compared to that in WT and mosaic mice, this should be reflected in the mis-expression of several activity-dependent iris muscle genes.

To quantify the expression of iris genes, we used bulk RNA sequencing. Of the ~13,000 genes detected, over 45% of these genes were dysregulated in iris muscle from α3 KO mice compared to those in wildtype iris muscle: 3520 genes were down-regulated at least 1.5-fold, 2757 genes were up-regulated, and the remaining 7100 were not statistically different (**Table S1**). These results indicate that in α3 KO mice almost half the iris muscle genes are dysregulated, which is consistent with long-term iris muscle inactivity and the absence of pupillary reflexes in these mice. On the other hand, fewer than 6% of the genes in iris muscle from mosaic mice were expressed at significantly different levels from those in wildtype iris (**Figure 4K, Table S1**), suggesting that the activity of iris muscles in mosaic mice appears generally normal.

Interestingly, we detected a significant increase in RNA levels for NGF, CNTF and neurturin in iris muscle from α3 KO mice but not from mosaic mice (**Table S2**); the RNA levels for other sympathetic nerve growth factors were not statistically different among iris muscle from wild-type, α3 KO, or mosaic mice. Nor were there differences in RNA levels for known target-derived factors, such as the BMPs or TGF beta family members. Exceptions were: BMP3, BMP receptors 1a and 2, TGF beta 2, and TGF beta receptor; the RNA levels for these five were 2-3-fold lower in α3 KO iris muscle than in mosaic or wild-type iris (**Table S2**).

### Influence of long-term inactivity on gene expression profile in SCG neurons

Gene expression profiles underly both cell-type identity and cellular activities; however, distinguishing among these various influences is often difficult. Our results indicate that X^RFP^ and X^α3^ neurons in mosaic SCG and wild-type SCG neurons are phenotypically similar in the extension of dendrites, the activity of targets they innervate, and in the innervation by preganglionic axons. However, X^RFP^ neurons do not receive synaptic transmission and preganglionic synapses on X^RFP^ neurons are silent. Therefore, a comparison of gene expression profiles in X^RFP^ neurons to those in X^α3^ and wild-type SCG neurons should shed some light on how synaptic transmission influences these gene expression profiles. Similarly, both X^RFP^ neurons and α3 KO SCG neurons lack synaptic transmission, yet differ in the activity of their targets. A comparison of gene expression profiles in X^RFP^ neurons to those in α3 KO SCG neurons should shed some light on how target activity influences these gene expression profiles. Therefore, we used singlecell RNA sequencing (scRNAseq) techniques to determine gene expression profiles of these different neuronal populations.

### Gene expression profiles in wild-type SCG neurons

First, we generated single-cell gene expression datasets from one-month-old SCG in wild-type mice (see Methods and Materials). Briefly, we pooled SCG from five to seven mice and dissociated them into a single-cell suspension (~9,000–10,000 cells were loaded and ~6,000–7,000 cells were recovered, a recovery rate of ~60–70%, consistent with droplet-based technology (Zhang et al., 2019)). The single-cell suspension contained both sympathetic neurons and ganglionic non-neuronal cells. We filtered out cells with fewer than 200 transcripts and genes that were expressed in fewer than 3 cells, using the R package, Seurat (Butler et al., 2018; Stuart et al., 2019). We then isolated the neuronal population with a set of selection markers for adrenergic neurons: positive for Dbh (dopamine beta hydroxylase, an enzyme involved in noradrenaline synthesis), Ntrk1 (high-affinity receptor for NGF, TrkA), Slc6a2 (noradrenaline transporter, NET), and voltage-gated sodium (Nav) and calcium channel (Cav) genes, and negative for Notch1, 2 or 3, specific markers for the non-neuronal cells.

Of the cells expressing Dbh, Ntrk1, Slc6a2 and negative for Notch, over 95% also expressed the voltage-gated calcium channel genes, Cacna1a (Cav2.1) or Cacna1b (Cav2.2), genes that code for the α1 subunit of P/Q and N-type calcium channels, respectively, and voltage-gated sodium channel genes, Scn9a (Nav1.7) and/or Scn3a (Nav1.3): approximately 75% expressed Scn9a, either alone or together with Scn3a, while approximately 25% of the neurons expressed Scn3a alone (**Figure S2**).

We identified the 1000 most highly variable genes and used a series of unsupervised learning algorithms to cluster the neurons into subtypes based on the differential (increased and decreased) expression of these highly variable genes relative to the total population. This process identified 7 distinct clusters, each comprising neurons with similar gene expression profiles. We refer to these clusters as SCG_1_–SCG_7_ and visualized them with a uniform approximation and projection method (UMAP) (**Figure 5A; Figure S3**) (Becht et al., 2019). These data indicate the degree of similarity and heterogeneity among neurons in the SCG. Moreover, the number of neurons that make up each cluster was not equal, suggesting that the specification of neuronal identity does not occur randomly.

**Figure 5.**
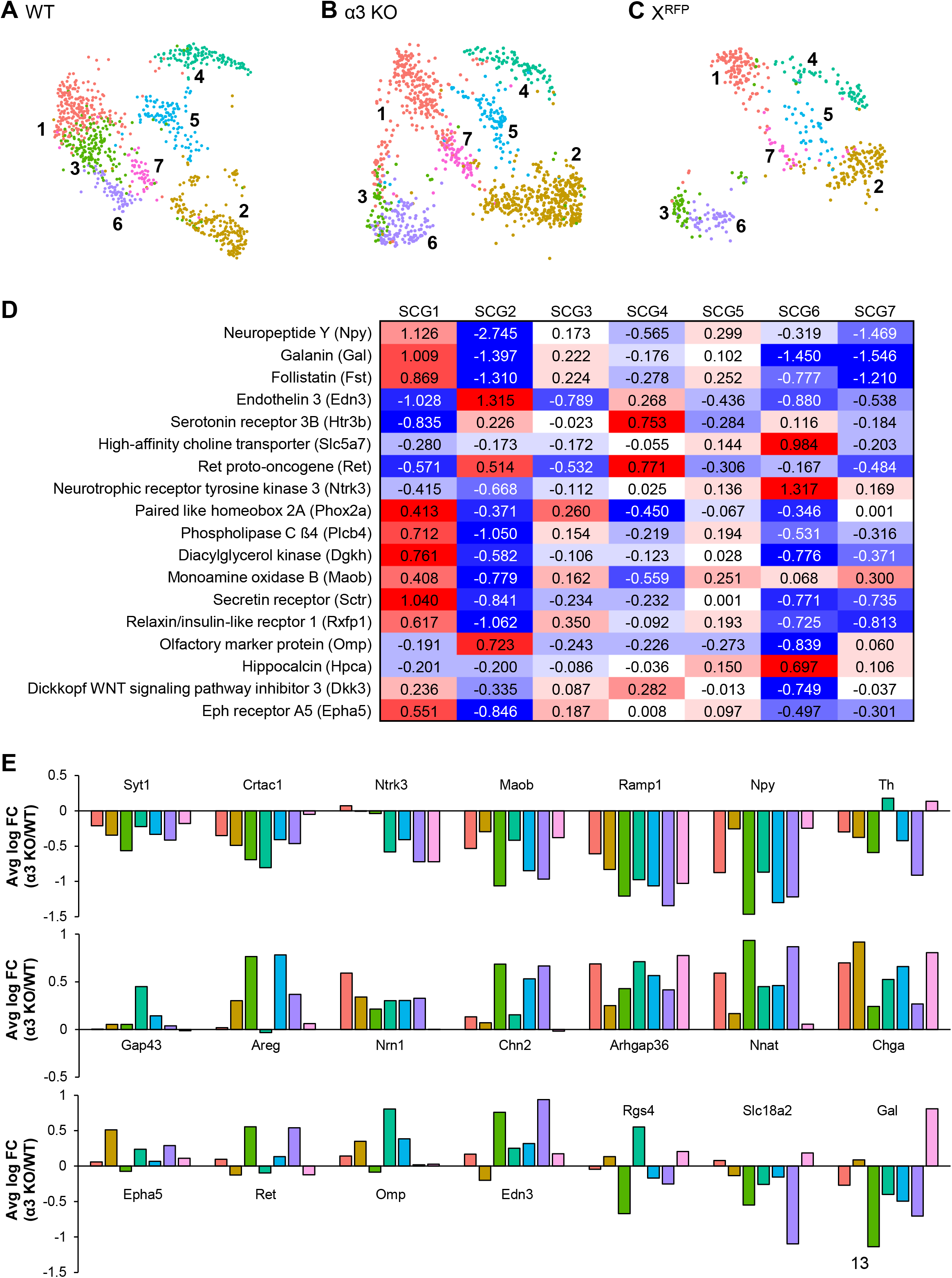
WT and α3 KO SCG neurons cluster into 7 subtypes. (**A-C**) UMAPs show clustering of neurons from (**A**) WT, (**B**) α3 KO and (**C**) mosaic SCG into 7 clusters. See also Figures S2, S3 and S4, and Tables S5 and S6. (**D**) Heat map showing the average ln(x+1) fold change of selected marker genes between neurons of a particular SCG clusters compared to the average expression in all other clusters. Each row was scaled independently. See also Table S3. (**E**) Subtypes of SCG neurons respond differently to changes in postsynaptic activity. Graphs show ln(x+1) fold change (α3 KO/WT) in each cluster for selected genes that were differentially expressed between WT and α3 KO SCG neurons. Negative fold change indicates downregulation in α3 KO neurons. The color of each bar refers to the SCG cluster shown in A. See also Table S4.

The marker genes that define each subtype are listed in **Table S3**. Among the marker genes are those that encode proteins known to be present in subpopulations of sympathetic neurons, including: NPY, 5HT_3_ receptors, galanin, TH, and VAChT, a cholinergic marker, CHT (high affinity choline transporter), Ret, Ntrk3, Phox2a, phospholipase C β4, diacylglycerol kinase, and monoamine oxidase b (**Figure 5D**). Immunohistochemistry or *in situ* hybridization experiments demonstrate that many of these “marker genes” are expressed in different sub-populations of rodent sympathetic neurons (Klimaschewski, Kummer, and Heym, 1996; Masliukov and Timmermans, 2004; Masliukov et al., 2012; Schütz et.al., 2008; Schotzinger and Landis, 1990). Significantly, no SCG cluster was defined exclusively by a single marker gene. Moreover, there were several marker genes whose expression and function has not been previously documented in sympathetic neurons; these include: the secretin receptor, the relaxin/insulin-like family receptor 1, olfactory marker protein, hippocalcin, Dickkopf-3, and EPH receptor 5A (**Figure 5D**).

### Gene expression profiles in SCG neurons from α3 KO mice

Next, we quantified gene expression profiles in SCG neurons from α3 KO mice and compared them to those in age-matched wild-type SCG neurons. To isolate the neuronal population, we used the same set of selection markers that we used for adrenergic neurons in wild-type SCG. We found no statistical differences in the expression of voltage-gated sodium and calcium channel genes between these α3 KO sympathetic neurons and those in wild-type SCG (**Figure S2**), indicating that the expression of these ion channel genes is not regulated by synaptic transmission. Next, we used a classifier, based on the dataset obtained from wild-type SCG neurons, to sort the α3 KO neuronal population into clusters corresponding to those in wild-type SCG (prediction score, 0.5). Over 95% of the 1115 neurons from α3 KO SCG clustered into the same 7 neuronal subtypes (**Figure 5B; Figure S3**). These data indicate that sympathetic neurons developing without synaptic transmission continued to mature into the same seven subtypes as wild-type neurons; however, more α3 KO SCG neurons were classified as SCG2 and SCG∈, than wild-type SCG neurons, and roughly 3-fold less as SCG3 (**Figure S3**).

Between wild-type and α3 KO SCG neurons, we identified 243 genes that were differentially expressed in at least one cluster (**Table S4**). SCG_2_ had the greatest number of differentially expressed genes (160 genes), whereas SCG7 had the lowest number (20 genes). In the absence of long-term synaptic transmission, many genes were either up-regulated or down-regulated in all clusters, including some genes known to be involved in refinement of presynaptic inputs or dendritic growth on CNS neurons (Leslie and Nedivi, 2011; Riccomagno and Kolodkin, 2015). Compared to wild-type SCG neurons, several genes, including those for neuropeptides, signaling molecules, and growth factor receptors were either up-regulated or down-regulated in some α3 KO SCG subtypes but unchanged in others (**Figure 5E**), suggesting that these genes are regulated by synaptic activity, however, this regulation depends on SCG subtype.

A particularly striking example is the expression of two markers for cholinergic neurons: CHT, the high-affinity choline transporter, and VAChT, the vesicular acetylcholine transporter. SCG_3_ neurons responded to the long-term absence of synaptic transmission by increasing the expression of CHT 40fold (α3 KO SCG3 versus wild-type SCG3) (**Figures 6A, B and G**), and VAChT by 90-fold (**Figures 6D, E and I**). We also observed a significant increase in these cholinergic markers in α3 KO SCG_6_ neurons. Moreover, there was a corresponding decrease in the expression of the adrenergic marker genes, VMAT2 and TH (**Figure 6K**). These data indicate that synaptic transmission in developing sympathetic neurons act to repress the expression of cholinergic genes.

**Figure 6.**
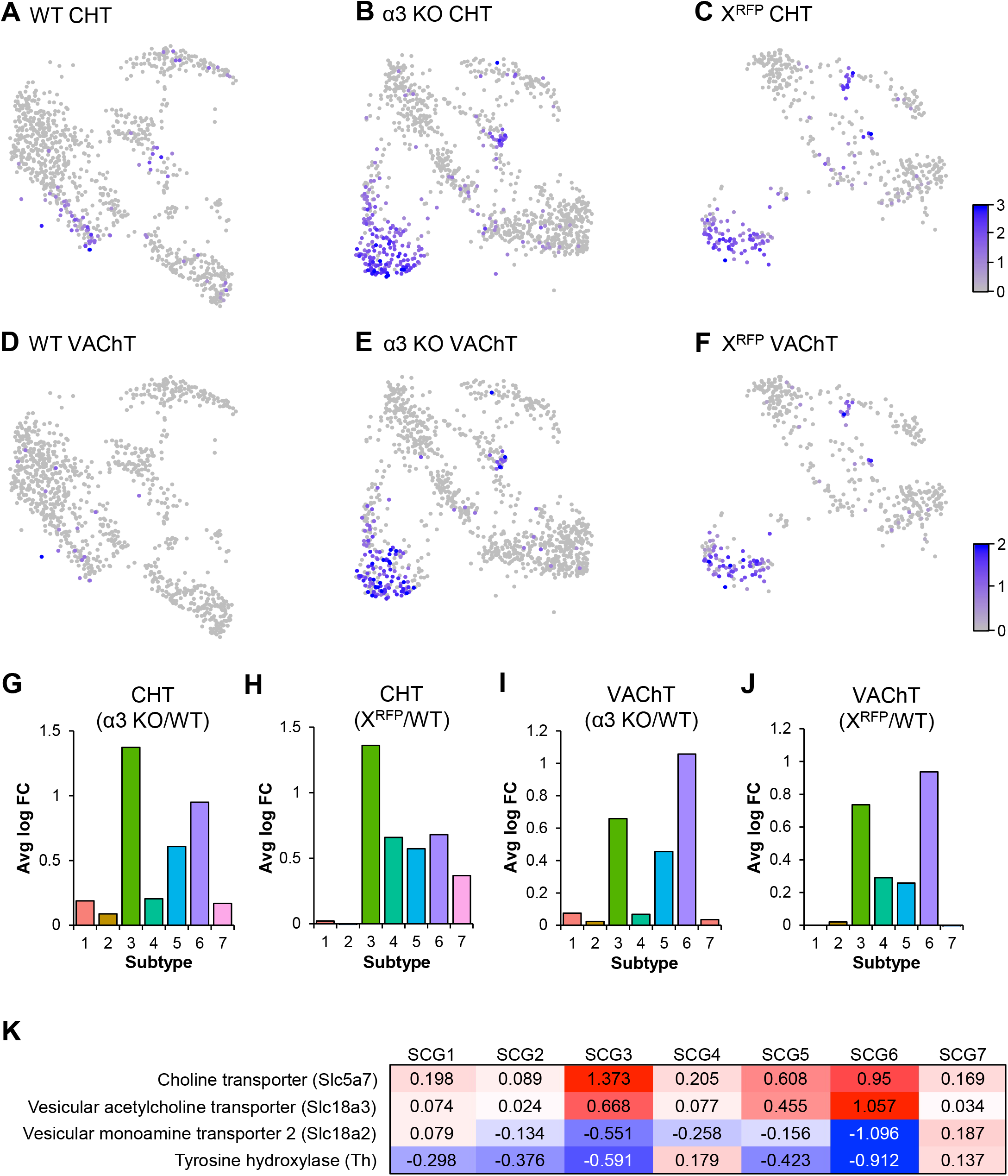
Cholinergic genes are expressed at higher levels in α3 KO and X^RFP^ SCG neurons than in WT SCG neurons. (**A-C**) UMAPs indicating expression level of CHT1 (Slc5a7) per neuron and their distribution between subtypes in (**A**) WT neurons, (**B**) α3 KO neurons, and (**C**) X^RFP^ neurons. (**D-F**) UMAPs indicating expression level of VAChT (Slc18a3) per neuron and their distribution between subtypes in (**D**) WT neurons, (**E**) α3 KO neurons, and (**F**) XRFP neurons. (**G, H**) Graphs show ln(x+1) fold change in each subtype for CHT1 (Slc5a7) between corresponding clusters in (**G**) WT and α3 KO neurons and (**H**) WT and X^RFP^ neurons. (**I, J**) Graphs show ln(x+1) fold change in each subtype for VAChT (Slc18a3) between corresponding clusters in (**J**) WT and α3 KO neurons and (**K**) WT and X^RFP^ neurons (**K**) Heat map showing the average ln(x+1) fold change of CHT1 (Slc5a7), VAChT (Slc18a3), VMAT2 (Slc18a2), and TH genes between neurons of a particular SCG clusters compared to the average expression in all other clusters. Each row was scaled independently. See also Table S3.

Relevantly, synaptic activity in the SCG had little influence on the genes expressed by non-neuronal cells. We identified 2409 non-neuronal cells in wild-type SCG and 2278 in α3 KO SCG. These non-neuronal cells clustered into 8 different subtypes (**Figure S4**); the marker genes for each cluster suggested that clusters 1, 2 and 4 appear to be ganglionic satellite cells, cluster 2 appears to be cells that make up the surrounding capsule, and cluster 6 appears to be mainly immune cells (**Table S5**). In comparing nonneuronal cells from wild-type SCG to those from α3 KO SCG, we identified only 22 genes that were differentially expressed by at least 1.25-fold (**Table S6**). The majority of these were housekeeping genes, and the expression of only 2 genes were different by more than 1.75-fold. These results indicate that synaptic activity in the SCG has little influence of the genes expressed by non-neuronal cells. None of these 22 genes have been implicated in the differentiation of developing sympathetic neurons.

These results with SCG neurons in α3 KO mice indicate that the long-term absence of synaptic transmission has a profound effect on the gene expression profiles of single SCG neurons. In α3 KO mice, however, not only is synaptic transmission absent at preganglionic synapses on SCG neurons, but the activity targets of SCG neurons is reduced significantly; both factors likely contribute to the expression of genes by SCG neurons. To distinguish between the influence of postsynaptic activity and target activity, we investigated gene expression profiles on X^RFP^ neurons in mosaic SCG. Like SCG neurons in α3 KO mice, X^RFP^ neurons do not receive synaptic transmission, however, the targets of these neurons appear active.

### Gene expression profiles in X^RFP^ neurons

To quantify gene expression profiles in X^RFP^ and X^α3^ neurons, we generated single-cell gene expression datasets from one-month-old mosaic SCG. The neuronal population was isolated with the same a set of selection markers for adrenergic neurons in wild-type SCG, and then we clustered the neurons using the classifier based on the wild-type data set. We recovered 1708 neurons from mosaic SCG and over 95% clustered into the same seven subtypes as wild-type SCG neurons.

X^RFP^ SCG3 and X^RFP^ SCG6 neurons had a significant increase in the number of neurons expressing CHT (**Figures 6C and H**) and VAChT (**Figures 6F and J**), as well as an increase in CHT and VAChT mRNA levels per neuron compared to wild-type SCG_3_ and SCG_6_ and a corresponding decrease in the expression of the adrenergic marker genes, VMAT2 and TH (**Figure 6K**). On the other hand, in X^α3^ neurons the expression of CHT and VAChT mRNA was not significantly different from that in neurons from wild-type SCG. These data reinforce the idea that synaptic transmission in developing sympathetic neurons acts to repress cholinergic genes.

Of the 243 differentially expressed genes in α3 KO SCG, X^RFP^ neurons expressed 156 (65%) of these at levels comparable to those in neurons α3 KO SCG, including CHT and VAChT (**Table S4**). These data indicate that these 156 genes are directly regulated by signals immediate downstream of postsynaptic activity.

On the other hand, X^RFP^ neurons expressed 87 of 243 (35%) genes at levels comparable to SCG neurons in wild-type mice (**Table S4**). Since X^RFP^ neurons do not receive synaptic transmission, these 87 genes are not regulated directly by postsynaptic transmission (or the resultant elevations in intracellular calcium), but by some process, possibly related to the activity of the target.

## Discussion

We generated mice with mosaic sympathetic ganglia composed of synaptically-silent X^RFP^ neurons that developed side-by-side with synaptically-active X^α3^ neurons, and we used these mosaic ganglia to investigate: 1) whether postsynaptic activity is required to initiate refinement of converging preganglionic inputs; and 2) how the long-term absence of synaptic activity influences gene expression profiles.

Our results show that preganglionic inputs to developing X^RFP^ neurons in mosaic SCG refine without synaptic transmission. This indicates that refinement of converging inputs to X^RFP^ neurons cannot be initiated by factors immediately downstream of postsynaptic activity and some other factor(s) must be involved. These results contradict widely-held views that refinement or pruning of inputs requires synaptic activity (Cohen-Cory, 2002; Kano and Hashimoto, 2009; Katz and Shatz, 1996; Lichtman and Colman, 2000; Zhang and Poo, 2001; Faust, Gunner and Schafer, 2021).

Previously, we showed that preganglionic axons innervating SCG neurons in α3 KO mice do not refine without synaptic activity (Chong et al., 2018), in contrast to what we show for inputs innervating X^RFP^ neurons. Since synaptic transmission is absent on both α3 KO SCG neurons and X^RFP^ neurons, postsynaptic activity does not initiate refinement. Instead, other differences between these two models presumably explain why inputs to X^RFP^ neurons refine whereas those that contact α3 KO SCG do not.

It seems unlikely that genetic manipulations to create these two mouse strains account for the difference. Both strains have a deletion in α3 on chromosome 9; however, mosaic mice have an additional modification on the X chromosome to express α3 cDNA or RFP. As a result of X-chromosome inactivation, in female mosaic mice, 50% of the neurons in the SCG express RFP (X^RFP^ neurons); these X^RFP^ neurons do not have functional nAChRs, similar to those in α3 KO SCG. Consequently, preganglionic inputs fail to evoke postsynaptic depolarizations on both α3 KO SCG neurons and X^RFP^ neurons.

A number of reports indicate that astrocytes and microglia have a role in refining neural connections (Chung et al., 2013; Neniskyte and Gross, 2017; Riccomagno and Kolodkin, 2015; Wilton et al., 2019; Faust, Gunner and Schafer, 2021). However, it seems doubtful that this occurs in sympathetic ganglia: unlike astrocytes in the CNS, ganglionic satellite cells have a one-to-one relationship with sympathetic neurons, which make it improbable that satellite cell signaling at active synapses on X^α3^ neurons could affect preganglionic connections on distant X^RFP^ neurons. More compelling, we found no significant difference between gene-expression profiles in satellite cells from WT SCG and those from α3 KO SCG (**Figure S5**) (**old S4**). Given this, instructing signal(s) for refinement of preganglionic inputs presumably come from elsewhere.

Another possibility is that a local activity-dependent factor emanating from X^α3^ neurons causes indiscriminate elimination of inputs to X^RFP^ neurons. However, this seems unlikely because elimination of preganglionic inputs is not a random process, but one that is usually thought to improve functional circuitry. The pupillary reflex is a good example. For pupillary reflexes to function independently, for instance while attending to a visual scene, SCG neurons innervating the iris require a unique set of preganglionic inputs to ensure that the nervous system can exert specific control over iris muscle without affecting other parts of the sympathetic nervous system (Neuhuber and Schrödl, 2011). Iris-specific neurons make up only ~0.5% of the total number of neurons in the SCG, and for the reflex to function properly, a specific set of preganglionic axons innervate iris-specific SCG neurons.

There is a striking difference in the activity of targets of sympathetic neurons between mosaic mice and α3 KO mice. We found that in α3 KO mice the activity of sympathetic targets are reduced considerably, or absent, whereas in mosaic SCG, the neighboring X^α3^ neurons continuously activate targets that they share with X^RFP^ neurons. These results highlight the importance of the target activity in initiating refinement of upstream inputs, as opposed to signals generated locally at synapses on postsynaptic neurons. A recent study on parasympathetic neurons concluded similarly that target innervation dictates the connections of central synapses (Sheu et al., 2017).

There are two likely ways that target activity could initiate the refinement of converging inputs to upstream neurons. One mechanism is through afferent feedback from the target and highlights the importance of overall circuit activity. Using pupillary reflexes as an example, preganglionic inputs to ~35 SCG control pupil diameter through iris muscle contraction. In mosaic mice, pupillary reflexes are largely intact because synapses between preganglionic terminals and X^α3^ neurons are functional. In this case, preganglionic neurons innervating iris-specific sympathetic neurons receive afferent feedback from the iris to modulate their firing rate and allowing them to out-compete other preganglionic neurons for iris-specific SCG neurons, even in the absence of postsynaptic activity. In α3 KO mice, on the other hand, iris muscle is mostly quiescent because all synapses between preganglionic terminals and postsynaptic sympathetic neurons are functional disrupted. Consequently, in α3 KO mice, pupillary reflexes are absent and afferent activity from iris muscle is significantly reduced. Accordingly, there is no afferent signals to allow iris-specific preganglionic neurons to out-compete other preganglionic neurons for iris-specific SCG neurons, and therefore refinement does not occur.

An alternative explanation for how target activity could initiate refinement, not necessarily exclusive from the first, is through retrograde factors released by targets in an activity-dependent manner. For example, over ~45% of the 13,000 genes detected in iris muscle from α3 KO mice were dysregulated, whereas less than 6% were dysregulated in iris from mosaics. Target-derived signals, such as NGF and BDNF, acting in a retrograde manner, play critical roles in the development and innervation of neurons (Choo et al., 2017; Cohen-Cory et al., 2010; Davies, 2009; Deppmann et al., 2008; Harrington and Ginty, 2013; Huang and Reichardt, 2001; Purves et al., 1988; da Silva and Wang, 2011; Singh et al., 2008; Snider and Lichtman, 1996; Zweifel, Kuruvilla and Ginty, 2005). Therefore, rather than local signals at synapses on postsynaptic neurons, this model proposes that the refinement of converging preganglionic axons is initiated by activity-dependent retrograde signals that originate from the targets of sympathetic nerves.

Relatedly, the density of sympathetic axons innervating iris muscle in α3 KO mice was 50% less than that in either wild-type or mosaic mice. It is not that the axons of synaptically-silent neurons grew less well because the axonal density of synaptically-silent X^RFP^ neurons innervating the iris in mosaic mice was similar to that of synaptically-active X^α3^ neurons. Instead, these results suggest that the activity of the target plays a major role in determining the target’s innervation density.

The density of target innervation has been shown to be regulated by retrograde growth factors (Davies, 2009; Huang and Reichardt, 2001), and mRNA levels for most of these growth factors in iris muscle from α3 KO mice were not statistically different than those in wild-type iris. Interestingly, we detected a significant elevation in neurturin, NGF, and CNTF mRNA in iris muscle from α3 KO mice, suggesting that the expression of these genes is regulated by muscle activity. Surprisingly, however, the up-regulation of these growth factor genes negatively correlated with the growth of sympathetic axons, suggesting that these growth factors by themselves are not sufficient to increase the growth of axons at the target. Presumably other factors, possibly specific to each target, are required to work in conjunction with these growth factors. For example, it was recently shown that S100b specifically promotes sympathetic innervation of brown adipose tissue (Zeng et al., 2019).

We find that dendritic outgrowth on X^RFP^ neurons was not statistically different from those on X^α3^ neurons in mosaic SCG or from age-matched wild-type SCG neurons. This was unexpected because previous work indicated that activity promotes the growth of dendrites on sympathetic neurons (Miller and Kaplan, 2003; Snider, 1988; Voyvodic, 1989). Moreover, we showed recently that SCG neurons in α3 KO mice have a significant decrease in dendritic outgrowth (Chong et al., 2018). That synaptically-silent neurons extend normal dendrites is not unique to X^RFP^ neurons in mosaic SCG; a sparse population of synaptically-silent CA1 neurons have normal dendrites (Lu et al., 2013). While the refinement of converging inputs to these CA1 neurons was not measured, relevantly, the density of synapses on the dendrites was not statistically different from control.

Previously, we showed that α3 KO SCG neurons have decreased amounts of phosphorylated 4E-BP, a repressor of cap-dependent mRNA translation, and have a corresponding change in the level of several proteins (Chong et al., 2018). Yet, we find that X^RFP^ neurons have phosphorylated 4E-BP levels that are comparable to those in X^α3^ neurons and SCG neurons in wild-type mice. Since synaptic transmission is absent X^RFP^ neurons, the signaling events that phosphorylate 4E-BP in X^RFP^ neurons can not come from nerve-evoked depolarization. Several studies have demonstrated that target-derived retrograde signals transported to soma alter gene expression at the transcriptional and translational level (Harrington and Ginty, 2013; Kaplan and Miller, 2000; Laplante and Sabatini, 2012; Lipton and Sahin, 2014; Segal and Greenberg, 1996; Tasdemir-Yilmaz and Segal, 2016). Therefore, it is conceivable that activity-dependent retrograde signaling from the target phosphorylated 4E-BP, and that poor target innervation in α3 KO mice may account for the decreased phosphorylated 4E-BP levels in SCG neurons in these mice.

We found a large up-regulation, both in the number of neurons and in average expression, in cholinergic marker genes, CHT and VAChT in α3 KO SCG, and a concomitant decrease in adrenergic marker genes, TH and VMAT2. Similarly, these cholinergic marker genes are up-regulated in X^RFP^ neurons but not in X^α3^ neurons in mosaic SCG. These results indicate that preganglionic nerve-evoked depolarization plays a critical role in maintaining an adrenergic phenotype in sympathetic neurons and are in line with those from other systems showing that synaptic activity influences the expression of neurotransmitters (Spitzer, 2012). Neonatal rodent SCG neurons developing in cell culture switch their transmitter phenotype from noradrenergic to cholinergic (Furshpan, Potter and Landis, 1979; Yamamori et al., 1989), however, consistent with a role for synaptic activity, this switch can be prevented by membrane depolarization (Walicke et al., 1977). In adult wild-type SCG *in vivo,* the majority of neurons are adrenergic and a small minority are cholinergic (Ernsberger and Rohrer, 1999; Schotzinger and Landis, 1990). Embryonically, however, sympathetic neuroblasts display a mixed noradrenergic and cholinergic phenotype, and transcriptional regulatory mechanisms and neurotrophic tyrosine kinase receptors, working through cross-regulatory interactions, determine whether a sympathetic precursor develops into an adrenergic or cholinergic neuroblast (Furlan et al., 2013). It is not clear whether synaptic activity engages this mechanism to prevent SCG neurons from acquiring cholinergic properties.

Using scRNAseq to analyze basal gene expression profiles, we show that SCG are composed of several subtypes of sympathetic neurons, similar to that of other sympathetic ganglia (Furlan et al., 2016). Interestingly, we found that the expression of several marker genes in SCG neurons has not been described previously and may underlie unsuspected functional aspects of these neurons. Equally relevant, we show that different subtypes respond differently to endogenous synaptic activity. For example, the up-regulation of cholinergic genes is particularly pronounced in SCG_3_ and SCG_6_, clusters that represent ~20% of the total neuronal population. This indicates that some subtypes are more affected by the absence of synaptic transmission than others with respect to the expression of cholinergic gene.

We found over 240 genes were differentially regulated in subtypes of α3 KO SCG compared to wild-type control neurons. This is consistent with reports that excitatory synaptic activity regulates the expression of immediate early genes that subsequently alter the expression of downstream target genes and influence neuronal development and innervation (Leslie and Nedivi, 2011; Yap and Greenberg, 2018).

On the other hand, comparing gene expression in α3 KO SCG to that in X^RFP^ neurons, we found more than 35% (87 genes) were dysregulated in α3 KO neurons but were expressed at near WT levels in at least one subtype of X^RFP^ neurons. Since preganglionic inputs do not evoke postsynaptic depolarizations on either α3 KO or X^RFP^ neurons, this indicates that the regulation of these ~87 genes is not mediated by synaptic EPSPs and related calcium influx. Rather, these genes must be regulated by other factors; the most likely are activity-dependent signals emanating from the targets.

In conclusion, our results from mosaic SCG demonstrate that presynaptic inputs refine in the absence of postsynaptic activity. In addition, we show that different SCG neuronal subtypes respond to the absence of synaptic activity during postnatal development. This difference likely depends on context, including the targets they innervate, and whether these targets are active.

## Materials and Methods

### Mice

To generate X^α3^X^RFP^ mosaic mice, α3 rat cDNA was ligated into a previously modified Gateway entry vector pENTR1a (Thermo Fisher Scientific, Waltham, MA) between the human ubiquitin C promoter (UbiC) and bovine growth hormone polyA site (Yurchenko et al., 2007). The pENTR1a entry vector was then recombined *in vitro* into the HPRT gateway destination vector (Thermo Fisher Scientific), which contained homology arms for the HPRT locus on the X chromosome (**Figure S1**). The recombined vector was electroporated into BK4 embryonic stem (ES) cells, which have a partial deletion in the HPRT gene. Successful homologous recombination inserted the UbiC-α3 construct and restored HPRT expression, which conferred resistance to 0.1mM hypoxanthine, 0.0004mM aminopterin, 0.016mM thymidine (HAT). HAT-resistant ES cells were injected into blastocysts that were transplanted into recipient females. Chimeric progeny that had germline expression of UbiC-α3 from the X chromosome were crossed with α3 KO mice and with X^RFP^ mice (Yurchenko et al., 2007) to generate α3 KO; X^α3^X^RFP^ mice. All genotyping was performed by PCR:

> Common UbiC FWD: 5’ – GCA GTG CAC CCG TAC CTT TGG GAG – 3’
>
> X^α3^ REV: 5’ – CTT AAA GAT GGC CGG CGG GAT CC – 3’
>
> X^RFP^ REV: 5’ – CGT AGG CCT TGG AGC CGT ACT GG – 3’

Mice with a deletion in the α3 nAChR subunit gene (α3 KO) were maintained on an outcrossed background because inbred C57BL/6J α3 KO mice die during the first week after birth (Krishnaswamy and Cooper, 2009). Briefly, inbred C57BL/6J α3+/− mice were mated to CD-1 WT mice and F1 α3+/− heterozygotes were used as breeders to produce α3 KO mice and WT littermates on a mixed C57BL/6J x CD-1 background. All genotyping was performed by PCR:

> Common FWD: 5’ – GTT ATG CAC GGG AAG CCA GGC TGG – 3’
>
> WT REV: 5’ – GAC TGT GAT GAC GAT GGA CAA GGT GAC – 3’
>
> α3 KO REV: 5’ – TGG CGC GAA GGG ACC ACC AAA GAA CGG – 3’

All procedures for animal handling were carried out according to the guidelines of the Canadian Council on Animal Care.

### Electrophysiological Recordings

SCG were acutely dissected in oxygenated Tyrode’s solution (pH 7.4) supplemented with glucose (5.6mM) and choline (0.01mM), pinned down securely with minutia pins on a Sylgard-coated petri dish, mounted to the stage of an upright confocal microscope (BX-61W, Olympus), and viewed through a 40X water-immersion objective (N.A. 0.8, Olympus). To record intracellularly from ganglion cells, 80–120mΩ glass microelectrodes (G150F-4; Warner Instruments, Hamden, CT) were made with a DMZ universal puller (Zeitz Instruments, Munich, Germany). Stable intracellular recordings were achieved with a high inertial precision microdrive (Inchworm 8200; EXFO, Quebec, Canada) attached to a micromanipulator (MPC-200/ROE-200; Sutter Instruments, Novato, CA) that drove the electrode through the ganglion at 4μm steps. The recording electrode was filled with 1M KAc. To confirm the identity of neurons in mosaic SCG of X^α3^X^RFP^ mice, 10mM Alexa Fluor 488 hydrazide (Thermo Fisher Scientific) in 200mM KCl was added to the electrode solution and neurons were imaged with confocal microscopy (Olympus). The recording electrode was connected with a silver chlorided wire to the head stage of an Axoclamp 2A amplifier (Axon Instruments, Union City, CA) used in current-clamp mode to deliver depolarizing or hyperpolarizing current pulses through the recording electrode. Ionic currents were filtered at 3kHz (low-pass cutoff) and 1Hz (high-pass cutoff) and digitized at 50kHz. Stimulation and data acquisition were performed with N-Clamp (Neuromatic, UK).

To measure the convergence of preganglionic axons innervating a sympathetic neuron, the preganglionic nerve was stimulated with increasing strength while holding the neuron at approximately –90mV to prevent EPSPs from triggering action potentials. In some experiments, the sodium channel blocker QX314 was also included in the recording electrode to prevent action potentials.

Analysis: All data analysis of electrophysiological recordings was performed offline using Igor Pro (WaveMetrics, Lake Oswego, OR). Increasing the strength of the stimulus to the preganglionic nerve activated axons of different threshold, which resulted in discrete jumps in the amplitude of the EPSPs. To isolate the average EPSP evoked by an axon and all axons of lower threshold, at least 10 traces were averaged for each discrete jump. The number of discrete jumps was used as an estimate of the number of axons innervating the neuron.

### Adenoviruses

Full-length α3 neuronal nAChR subunit cDNA was ligated into pAdTrack-synapsin 1 (Ad-α3/Syn), and replication-deficient viral vectors were generated (He et al., 1998) and titered in duplicate with Adeno-X Rapid Titer Kit, (Clontech Lab, Mountain View, CA; Krishnaswamy and Cooper, 2009). Mice were infected with Ad-α3/Syn adenovirus at a concentration of ~10^7^pfu/mL diluted in sterile 1X PBS. For P5–P6 mice, ~50μL was injected into the intraperitoneal cavity, and for P23–P26 mice, ~200μL was injected intravenously into the tail vein. Injections were performed using a 29G x 1/2” 1mL insulin syringe (Bencton Dickinson, Franklin Lakes, NJ).

### Imaging

Images were acquired on an upright confocal microscope (BX-61W, Olympus) with a 60X, N.A. 1.42 PlanApo N oil-immersion objective (Olympus) at a scan speed of 8μs/pixel and an image depth of 12 bits. Laser lines were activated sequentially to avoid bleed-through of signals. All image analysis was performed with FIJI/ImageJ (NIH, Bethesda, MD).

### Lipophilic tracer labelling for dendrite morphology

Lipophilic tracer 3,3’-Dioctadecyloxacarbocyanine perchlorate (DiO) (Thermo Fisher Scientific) was used to sparsely label a random subset of preganglionic axons and postsynaptic neurons in the SCG. Briefly, freshly dissected ganglia with intact pre- and postganglionic nerves were fixed in 1% PFA (pH 7.4) in 0.1M PB for 2 hours at room temperature, rinsed with 1X PBS and embedded in 3% agarose Type I-B (Sigma-Aldrich, St. Louis, MO) dissolved in 1X PBS. A scalpel blade was used to slice through the pre- and postganglionic nerves to expose a cross-sectional area of the nerve. Platinum wires were used to gently apply fine crystals of DiO to the postganglionic nerve to label sympathetic neurons and their dendritic structures. After labelling, ganglia were kept in the dark in 1X PBS for 5–6 days to allow for tracers to diffuse along lipid membranes. Excess DiO was removed and ganglia were sliced into 100μm sections with the Compresstome VF-200 (Precisionary Instruments Inc., Greenville, NC) using a solid zirconia ceramic injector blade (Cadence Inc., Staunton, VA). Sections were either mounted with Vectashield (Vector Laboratories, Burlingame, CA) and immediately imaged, or first processed with immunohistochemistry before mounting and imaging.

Analysis: Only neurons with complete dendritic arbours and an identifiable axon were analyzed. For neurons with a dendritic arbour that extended beyond one field of view, neighboring z-stacks were acquired and stitched together with XuvTools (Emmenlauer et al., 2009). To quantify the length and number of dendritic branches, neurons were reconstructed in 3-D and the Simple Neurite Tracer plugin (Longair et al., 2011) was used to trace dendrites. Primary dendrites were categorized as those directly leaving the cell body.

For representative images, neurons were tiled and DiO-labelled neurites of other neurons were removed from the field of view for clarity. On each plane of the z-stack, all DiO-labelled neurites that were not connected to the dendritic arbour of the neuron of interest, as determined by 3-D reconstruction and dendritic tracing, were removed for clarity. After removing non-connected neurites, the z-stack was used to produce a maximum intensity z-projection of the neuron. For illustration purposes, figures show the maximum intensity z-projections.

### Synaptic targeting on DiO labelled neurons

SCG were fixed and the postganglionic nerve was labelled with DiO and incubated in the dark for 5–6 days as described above. Labelled SCG were sliced into 100μm sections and incubated in blocking solution for 2 hours at room temperature [Blocking solution: 10% normal donkey serum (Millipore, Billerica, MA) and 0.3% Tween 20 (Fisher Scientific, Waltham, MA) in 1X PBS], in the primary antibody for 48 hours at 4°C [Primary antibody: Rabbit anti-VAChT (1:3000; Synaptic Systems)], and in the secondary antibody diluted in 10% normal donkey serum for 2 hours at room temperature [Secondary antibody: Alexa Fluor 647 goat anti-rabbit (1:500; Thermo Fisher Scientific)].

Analysis: To examine synaptic targeting, VAChT puncta located on a neuron of interest were identified on each plane of a z stack. To be counted as a synapse, VAChT puncta were: (i) colocalized with the DiO membrane label; (ii) at least 0.5μm in diameter; and (iii) spanned at least two optical slices (optical thickness was 0.45μm/slice). Synapses were categorized as being on the cell body or on the dendritic arbour. The density of synapses on the cell body was calculated by dividing the number of synapses on the cell body by the total surface area of the cell body. To estimate the surface area of the cell body, the circumference of the cell body on each plane was multiplied by the thickness of the optical slice and summed. To calculate the density of synapses on dendrites, the number of synapses on dendrites was divided by the total dendritic outgrowth (TDO). TDO was measured using 3-D reconstructed neurons and the Simple Neurite Tracer plugin as described above. For representative images, VAChT puncta that were not located on the neuron of interest were removed for clarity. To illustrate clearly the location of VAChT puncta, maximum intensity z-projections of DiO-labelled SCG neurons were thresholded and skeletonized, and the locations of VAChT puncta were indicated with red circles.

### Phosphorylated-4E-BP (P-4E-BP) immunohistochemistry

SCG were dissected and immediately fixed in 2% PFA (pH 7.4) in 0.1M PB with 5mM EGTA (to chelate Ca^2+^ ions released from intracellular stores by fixation) for 1 hour at room temperature. After fixation, ganglia were sliced into 100μm sections and incubated in blocking solution for 1 hour at room temperature [Blocking solution: 10% normal donkey serum, 0.3% Tween 20, and 0.05% Triton X-100 (Fisher Scientific) in 1X PBS], in primary antibodies for 48 hours at 4°C [Rabbit anti-P-4E-BP (1:600; Cell Signaling Technology, Danvers, MA), goat anti-MAP-1A (1:360; Santa Cruz Biotechnology, Dallas, TX)], and in secondary antibodies diluted in 10% normal donkey serum for 1 hour at room temperature [Secondary antibodies: TRITC donkey anti-rabbit (1:500; Jackson ImmunoResearch Laboratories), Alexa Fluor 647 donkey anti-goat (1:500; Thermo Fisher Scientific)].

Analysis: All image acquisition parameters (HV, gain, offset, laser power) were kept constant between samples. Z stacks of 20 images were used to generate a summed z-projection and regions of interest (ROI), each consisting of one neuronal cell body, excluding the nucleus, were selected from the MAP-1A channel. Average fluorescence intensity for each ROI was measured from the MAP-1A channel, and ROI were transferred to the P-4E-BP1 channel to measure the corresponding fluorescence intensity. Figures show maximum intensity z-projections.

### Sympathetic innervation of the iris

Irises were dissected from albino mice and fixed with 2% PFA (pH 6.0) in 0.1M PB for 10 minutes at room temperature. To achieve a consistent state of contraction between samples, carbachol was added to the fixative for a final concentration of 500μM. During fixation, irises were left attached to the cornea to maintain the structure and carefully separated from the cornea with a scalpel blade after fixation. Irises were incubated in blocking solution for 1 hour at room temperature [Blocking solution: 10% normal donkey serum, 0.3% Tween 20, and 0.1% Triton X-100 in 1X PBS], in primary antibodies for 24 hours at 4°C [Primary antibodies: Mouse anti-tyrosine hydroxylase (TH) clone LNC1 (1:500; Millipore) and rabbit anti-vesicular monoamine transporter 2 (1:1000; Phoenix Pharmaceuticals, Burlingame, CA)], and in secondary antibodies diluted in 10% normal donkey serum for 1 hour at room temperature [Secondary antibodies: Alexa Fluor 405 goat anti-rabbit (1:500; Thermo Fisher Scientific) and Alexa Fluor 647 goat anti-mouse IgG1 (1:500; Thermo Fisher Scientific)].

Analysis: All image acquisition parameters (HV, gain, offset, laser power) were kept constant between samples. Images were taken at the radial muscles halfway between the inner (pupil) and outer (attachment to sclera) edge of the iris. Z-stacks were split into separate channels for TH and VMAT2, and the same threshold values for each channel were applied to all samples. On each channel, the mean fluorescence intensity of the pixels above threshold were recorded. Innervation density was expressed in two ways: (1) TH-positive area over a given area, and (2) number of VMAT2 puncta in a given area. To calculate the TH innervation density, a maximum intensity z-projection was generated from the TH channel, thresholded, and the TH area was divided by the total area. To calculate the VMAT2 innervation density, a maximum intensity z-projection was generated from the VMAT2 channel, single pixel noise was removed with the “Despeckle” filter and the “Find maxima” filter was used to automatically count puncta (noise=700), and the number of puncta was then divided by the total area. To calculate the number of VMAT2 puncta along TH-positive fibers, a maximum intensity z-projection of the TH channel was thresholded, skeletonized, and total fiber length was measured. Figures show a maximum intensity z-projection for each channel.

To determine the ratio between X^α3^ axons and X^RFP^ axons in mosaic mice, subsets of 3 slices were summed through a maximum intensity z-projection and individual axon fibers were identified on the TH channel. The lengths and areas of the axons were measured, and the number of VMAT2 puncta along the axon was counted. Each axon was then overlaid onto the RFP channel, and categorized as either RFP-positive, RFP-negative, or undetermined.

### Retrograde labelling with cholera toxin subunit B (CTB)

P28 mice were anaesthetized with isoflorane and the cornea was punctured with a 29G hypodermic needle (Bencton Dickinson) to allow for outflow of aqueous humour. Using the same puncture site, ~2μL of cholera toxin subunit B conjugated to Alexa Fluor 488 (CTB-488; Thermo Fisher Scientific) was injected into the anterior chamber of the eye using a 33G x 1/2” TSK SteriJect hypodermic needle (Air-Tite Products, Virginia Beach, VA) fitted onto a 25μL Gastight Hamilton syringe (Hamilton Company, Reno, NV). SCG were dissected after ~4 days, fixed in 1% PFA in 0.1M PB for 1 hour at room temperature and sliced into 100μm sections for imaging.

Analysis: All labelled neurons were intensely fluorescent and easily identified: mean intensity fluorescence (excluding the nucleus) was at least 10X greater than background fluorescence intensity. For mosaic SCG, labelled neurons were superimposed on the RFP channel and categorized as RFP-positive or RFP-negative. Low magnification figures for illustration purposes were obtained using a 10X air objective (N.A. 0.3, Olympus).

### Gene expression in the iris

Six irises were collected from 3 mice for each genotype (WT, α3 KO and mosaic mice). Irises were dissected in DNase/RNase-free 1X PBS (DNase/RNase-free 10X PBS from Thermo Fisher Scientific diluted in DNase/RNase-free H2O) and immediately flash frozen in liquid nitrogen. Total RNA was extracted in parallel using the RNeasy Mini Kit (Qiagen, Hilden, Germany). Frozen tissue was homogenized (Polytron PT 2100, Kinematica, Luzern, Switzerland) in buffer RLT with β-mercaptoethanol. Homogenates were processed according to the RNeasy Mini Kit protocol and included a DNase I (Qiagen) treatment step to avoid contamination from genomic DNA. RNA quantity and purity was assessed with a NanoDrop2000c (NanoDrop, Wilmington, DE; 260/230 and 260/280 values > 2.0) and RNA integrity was assessed with a Bioanalyzer (Agilent Technologies, Santa Clara, CA; RIN > 8.0). cDNA libraries were sequenced with an Illumina NovaSeq 6000 System (S4, PE100; Illumina) at 25M reads.

Analysis: Count data for each transcript were generated from paired-end FASTQ files with Kallisto v0.46.1 (Bray et al., 2016) using a prebuilt kallisto index from the Mus Musculus Ensembl v96 transcriptome and transcripts were summarized into genes and imported into R Studio (R Core Team, 2013; RStudio Team, 2015) with the tximport package (Soneson et al., 2016). Genes with low counts were filtered out (cpm >= 3), and differential analysis was performed with the EdgeR package (McCarthy et al., 2012; Robinson et al., 2010) in R using generalized linear models (GLMs) for non-normal distributions. Genes with a log2 fold change > 0.58496 (1.5X) at a p value < 0.05 were considered as differentially expressed.

### Ultrastructural studies

Mosaic SCG were dissected, fixed in 4% PFA (pH 7.4) in 0.1M PB for 1 hour at room temperature, sliced into 100μm sections and fixed for an additional 10 minutes at room temperature. Sections were rinsed with 0.1M PB, imaged with a 40X water-immersion objective (N.A. 0.8, Olympus) on an upright confocal microscope (Olympus). After imaging, sections were fixed again in 2% PFA/2% glutaraldehyde for 30 minutes at room temperature. After fixation, SCG were rinsed with 0.1M PB for 30 minutes and incubated in 1% osmium tetroxide/1.5% potassium ferricyanide in H_2_O for 1 hour at room temperature. Ganglia were rinsed briefly with H_2_O to remove osmium tetroxide and dehydrated in a graded series of ethanol concentrations from 30–100%, where each interval lasted 10 minutes, with the 100% ethanol step repeated 3 times. After dehydration, ganglia were incubated in 100% propylene oxide for 15 minutes twice and incubated in a propylene oxide:EMbed812 mixture at a ratio of 1:1, 1:2, 1:3 and pure EMbed812 for 1 hour each, and polymerized in EMbed812 at 60°C for 24 hours. Thin sections of ganglia were cut on an ultramicrotome, stained with 2% aqueous uranyl acetate and 3% lead citrate, and viewed with a Tecnai Spirit 120kV transmission electron microscope with Gatan Ultrascan 4000 4k x 4k CCD Camera System Model 895.

### α3 mRNA expression from X chromosome

RNA extraction and quantitative PCR (qPCR) were optimized, validated and performed in accordance with the MIQE guidelines (Bustin et al., 2009; Taylor et al., 2010). SCG from P28 WT, α3+/− heterozygote, and mosaic mice were dissected and immediately flash frozen in liquid nitrogen. Total RNA was extracted using the RNeasy Mini Kit (Qiagen). Frozen tissue was homogenized (Polytron PT 2100, Kinematica) in buffer RLT with β-mercaptoethanol. Homogenates were processed according to the RNeasy Mini Kit protocol and included a DNase I (Qiagen) treatment step to avoid contamination from genomic DNA. RNA quantity and purity were assessed with a NanoDrop2000c (NanoDrop) and RNA integrity was assessed by running 400ng on a gel. 260/230 and 260/280 values were consistently >2.0, and 28S and 18S rRNA bands were consistently clear and sharp, with the 28S band approximately twice as intense as the 18S band. Reverse transcription (iScript Reverse Transcription Supermix for RT-qPCR, Bio-Rad, Hercules, CA) was performed immediately after RNA extraction and used 200ng of RNA per reaction volume of 20μL. No reverse transcriptase controls were included for each reaction to test for contamination from genomic DNA. cDNA was stored at −20°C.

qPCR reactions were performed using SsoFast EvaGreen Supermix with Low ROX (Bio-Rad, Hercules, CA) on the Eco Real-Time PCR System (Illumina, San Diego, CA). Cycling parameters were as follows: UDG incubation 2mins at 50°C; Polymerase activation 30s at 95°C; PCR cycling (5s at 95°C, 15s at 60°C) for 40 cycles. All primers were 90-100% efficient at an annealing temperature of 60°C. Samples were run in duplicates and controls without reverse transcriptase and template consistently showed no amplification.

In α3 KO mice and in mosaic mice, the endogenous α3 nAChR subunit gene on chromosome 9 had a deletion in exon 5 that prevented the formation of functional α3 subunit protein. The forward primer was designed to bind within exon 4 and reverse primer to bind within the deleted region in exon 5, and therefore would not recognize any potential mRNA generated from the KO allele. In addition, intron-spanning primer pairs reduced amplification of residual genomic DNA. Standard curves were generated from an 8-point 4X serial dilution of cDNA to determine primer efficiencies. α3 mRNA levels were normalized to GAPDH expression.

α3 FWD: 5’ – GTG GAG TTC ATG CGA GTC CCT G – 3’ (in exon 4)

α3 REV: 5’ – TAA AGA TGG CCG GAG GGA TCC – 3’ (in exon 5)

GAPDH FWD: 5’ – CTG GCA TGG CCT TCC GTG TT – 3’

GAPDH REV: 5’ – TAC TTG GCA GGT TTC TCC AGG CG – 3’

### Single cell RNA sequencing (scRNAseq) of sympathetic ganglia

Single cell RNA libraries were generated using 10X Genomics droplet-based technology at the Genome Québec Innovation Centre (Montréal, Canada). SCG from five to seven P28 mice were pooled for each genotype (WT, α3 KO and mosaic mice). Freshly dissected SCG were incubated on a shaker in 1mg/mL collagenase (Sigma-Aldrich) and 5mg/mL bovine serum albumin (BSA; Sigma-Aldrich) in 1X HBSS (pH 7.4) for 30 minutes at 37°C, followed by 1mg/mL trypsin (Worthington Biochemical Corporation, Lakewood, NJ) in 1X HBSS (pH 7.4) for 1 hour at 37°C. To dissociate ganglia into a single cell suspension, SCG were gently titurated with a fire-polished pipette, washed with 10% horse serum and 1X HBSS, and filtered through a 30μm cell strainer (Miltenyi Biotec, Bergisch Gladbach, Germany). Cells were reconstituted in calcium- and magnesium-free 1X PBS containing 400μg/mL non-acetylated BSA at concentration of ~1000 cells/μL in 2mL LoBind Eppendorf tubes (Thermo Fisher Scientific). Cell viability was determined with a 0.4% trypan blue solution (Thermo Fisher Scientific) and was consistently >90%. Cell suspensions were kept on ice and loaded onto the 10X Loading Single Cell Chip within 30 minutes to 1 hour after preparation. For all samples, ~9,000–10,000 cells were loaded and ~6,000–7,000 cells were recovered for a recovery rate of ~60–70%, consistent with this technology ^36^. RNA molecules were captured, tagged with a cellular barcode and a unique molecular identifier barcode, and reversed transcribed to generate cDNA libraries for sequencing.

Analysis: cDNA libraries were amplified and multiplexed for sequencing with the Illumina HiSeq 4000 (Illumina) to generate read data. Read data were grouped based on barcodes and aligned using Cell Ranger software to a custom reference package consisting of the mouse genome version GRCm38/mm10 with the addition of rat α3 and mRFP1 cDNA.

Count matrices were analyzed using the Seurat package v3.0 (Butler et al., 2018; Stuart et al., 2019) in Rstudio. Cells that had fewer than 200 transcripts (less than 0. 08 and 0.4% of the cells expressed fewer than 200 genes), and genes that were expressed in fewer than 3 cells were filtered out. (WT: total = 5781 cells; after filtering, 5776; α3 KO: total = 5587 cells; after filtering = 5580; mosaic: total = 13043 cells; after filtering = 12788 cells.) To be considered a neuron, cells expressed Dbh, Ntrk1, Slc6a2, either Scn3a or Scn9a (or both), and either Cacna1a or Cacna1b (or both), and not Notch1, 2 or 3. To isolate the non-neuronal population, we selected cells that expressed Notch1, 2 or 3 but not Dbh, Slc6a2, Scn3a, Scn9a, Cacna1a, Cacna1b nor Npy. The fraction of mitochondrial counts per cell was consistently <0.2 and there were no outliers with high transcript counts that may represent doublets. The neuronal subset was normalized to the count depth per cell and transformed via a natural log-plus-one transformation (ln(x+1)). For clustering analysis, data were scaled to have zero mean and variance of 1. Scaled expression equally weighed high- and low-expressing genes for clustering analysis. The top 2000 highly variable genes were identified and used for preprocessing with principal components analysis (PCA). The WT dataset was clustered and used as a classifier to sort α3 KO neurons, X^α3^ neurons and X^RFP^ neurons into the same clusters identified in the WT population (prediction score > 0.5). Clusters were visualized using a uniform approximation and projection method (UMAP; Becht et al., 2019). To test for differential expression, a Wilcoxon rank sum test was used. Genes were considered to be differentially expressed if their average ln(x+1) fold change was ±0.25, with a Bonferroni adjusted p-value <0.05.

Of the differentially expressed genes between α3 KO SCG and WT SCG, we determined whether their levels in X^RFP^ neurons from mosaic SCG were comparable to levels in α3 KO SCG, suggesting they are directly regulated by postsynaptic activity, or whether they were comparable to those in WT SCG, suggesting that these genes are indirectly regulated by activity. Using the normalized average expression level, the difference was calculated for each differentially expressed gene between α3 KO SCG and WT SCG. Then, 25% of the difference was either added onto the α3 KO value if it was down-regulated compared to WT, or subtracted from the α3 KO value if it was up-regulated compared to WT. If the X^RFP^ average expression level surpassed this 25% threshold in the direction of the WT value, the gene was considered to be restored towards WT values.

### Statistical analysis

The number of samples (n) and p-values are reported in the figures and corresponding figure legends. In all figures, error bars represent ± s.e.m., *p<0.05, ***p<0.001. To test for statistical differences between two samples, unpaired two-tailed t tests assuming equal variance were used.

## Acknowledgments

We are grateful to Drs. A. Krishnaswamy, S. McFarlane, and E. Ruthazer for comments and suggestions on previous versions of this manuscript, and Brigitte Pié for excellent technical assistance. Single-cell RNA libraries were generated using 10X Genomics droplet-based technology and sequenced at the Genome Québec Innovation Centre in Montréal, Québec. This work was supported by a Canadian Institute for Health Research operating grant.

## Author Contributions

Y.C. and E.C. were responsible conceptualization, investigation, methodology, data collection and analysis, visualization, writing – original draft, and review and editing, E.C. was responsible for funding acquisition.

---

**Figure S1.**
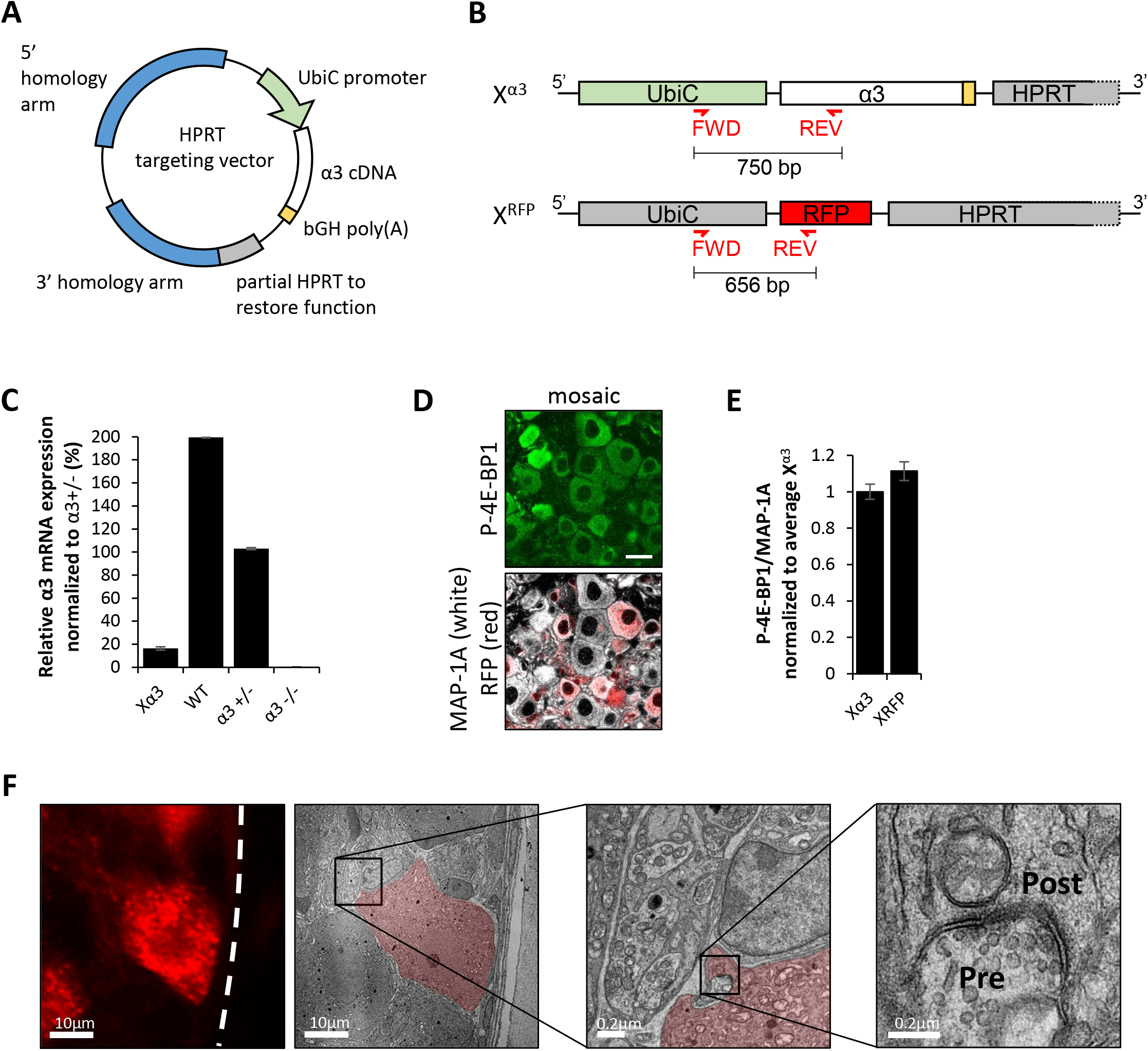
Design of the mosaic mouse model. (**A**) α3 rat cDNA (white) was ligated into a previously modified vector in between the human ubiquitin C promoter (UbiC; green) and bovine growth hormone polyA site (bGH poly(A); yellow) and recombined *in vitro* into the HPRT gateway destination vector to generate the targeting vector. Homology arms (blue) direct homologous recombination into the HPRT locus, and partial HPRT (grey) restores HPRT function in HPRT-deficient ES cells for resistance to HAT media (see Materials and Methods). (**B**) Diagram shows locations of α3 and mRFP1 genes between the human ubiquitin C promoter (UbiC) and HPRT gene on the X chromosome. Forward (FWD) and reverse (REV) primers used for genotyping are indicated in red with expected amplicon sizes, primer sequences can be found in Materials and Methods. (**C**) Relative α3 mRNA expression in SCG normalized to α3+/− SCG. For X^α3^, n=22 mice; for WT, n=3 mice, for α3+/−, n=3 mice; and for α3 KO, n=2 mice. (**D**) Confocal images showing immunostaining for P-4E-BP1 (green) and MAP-1A (white) in mosaic SCG (RFP expression in red) at P28. Scale bar, 20μm. (**E**) P-4E-BP1 mean fluorescence intensity per neuron normalized to MAP-1A in X^α3^ and X^RFP^ neurons at P1, P4 and P28. (**F**) Corresponding images of a X^RFP^ neuron viewed at the light microscopy level and at the ultrastructural level (red). Boxed areas were magnified to show presynaptic vesicles and postsynaptic densities.

**Figure S2.**
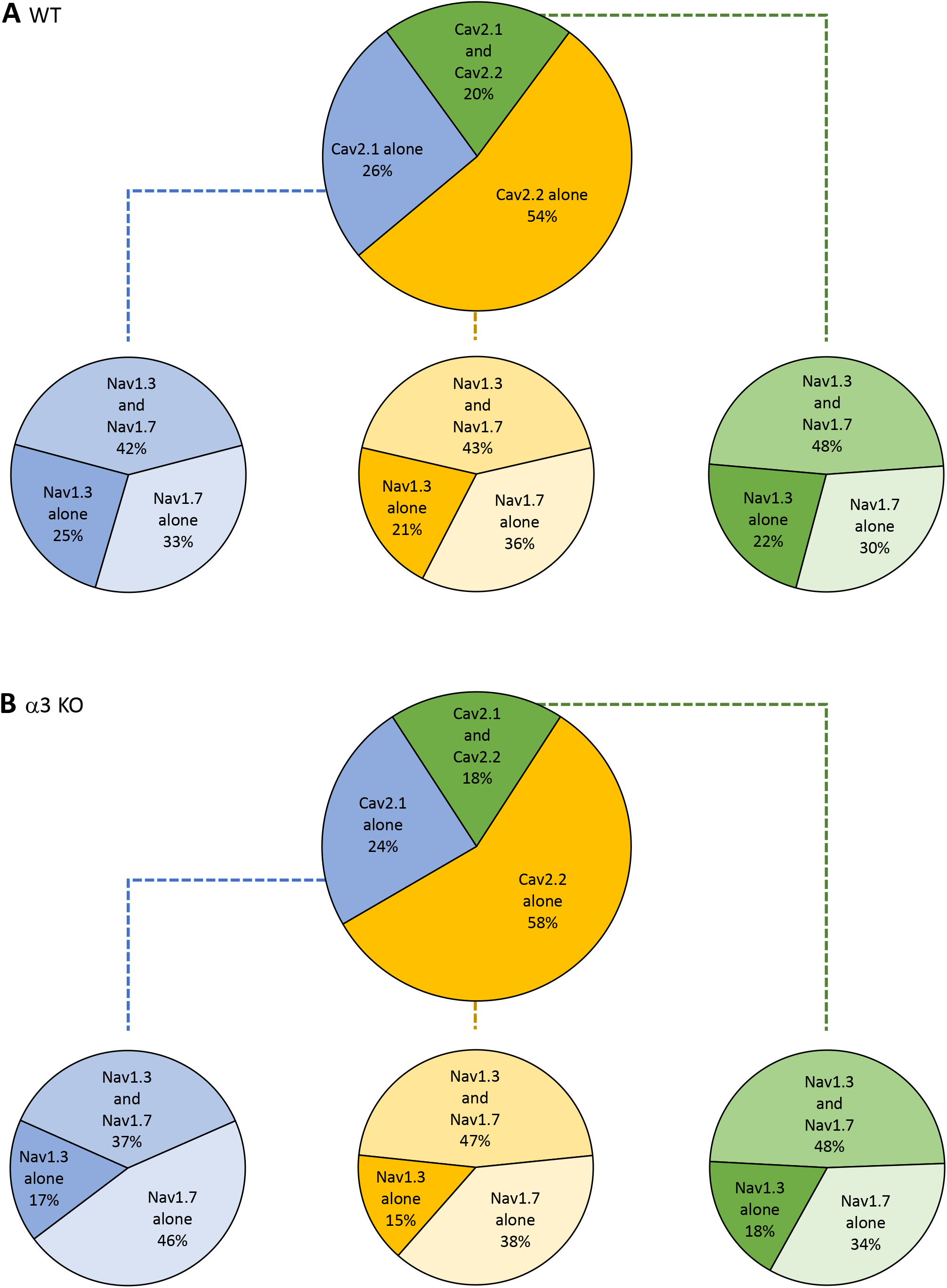
Expression of αi Ca_v_ subunit and α Na_v_ subunit genes in WT and α3 KO neurons. (**A-B**) Top: Distribution of neurons that expressed Cacna1a without Cacna1b (Cav2.1; blue), Cacna1b without Cacna1a (Cav2.2; yellow), or both (green) in (**A**) WT SCG and (**B**) α3 KO SCG. Bottom: Each of the three Cav groups were further broken down according to the distribution of neurons that expressed Scn3a without Scn9a (Nav1.3), Scn9a without Scn3a (Nav1.7), or both.

**Figure S3.**
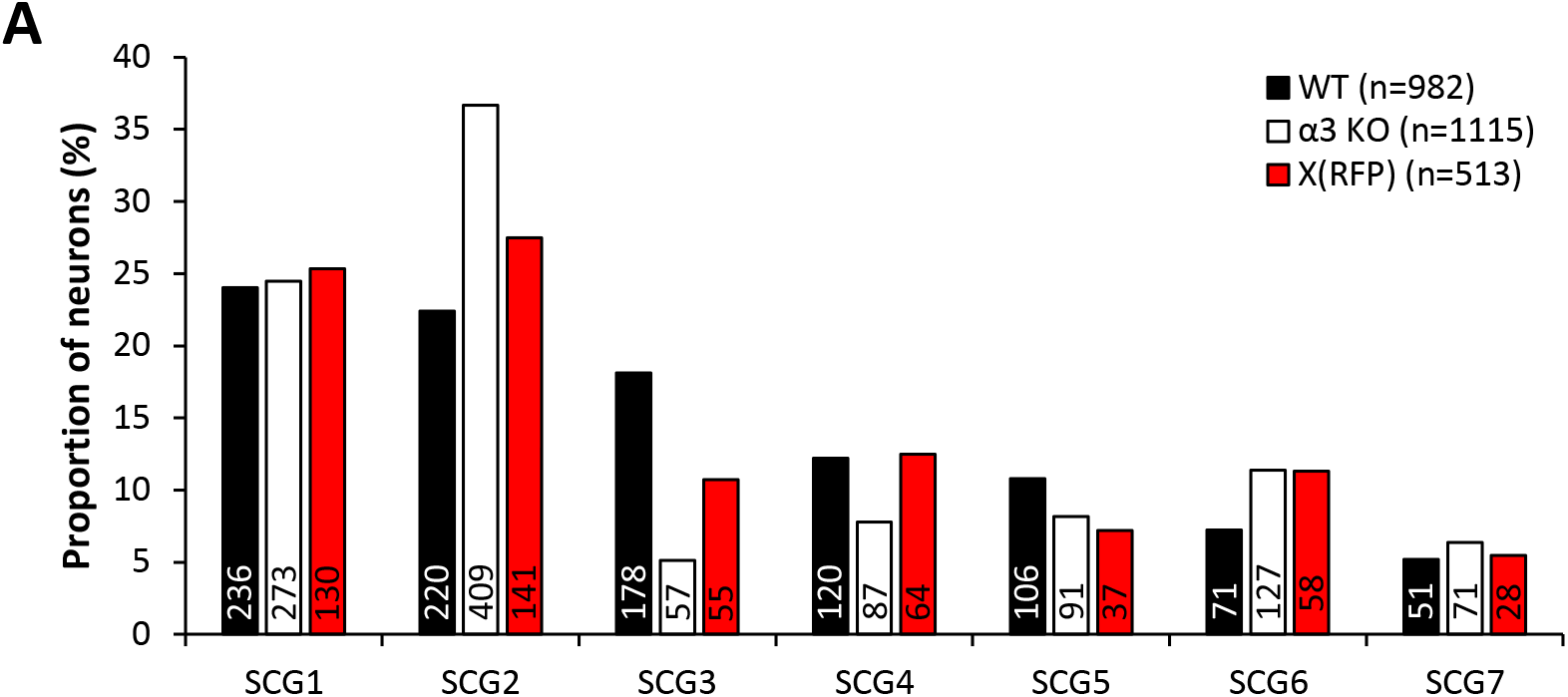
SCG neurons were clustered into 7 subtypes. The proportion of WT (black), α3 KO (white) and X^RFP^ (red) SCG neurons categorized into each subtype.

**Figure S4.**
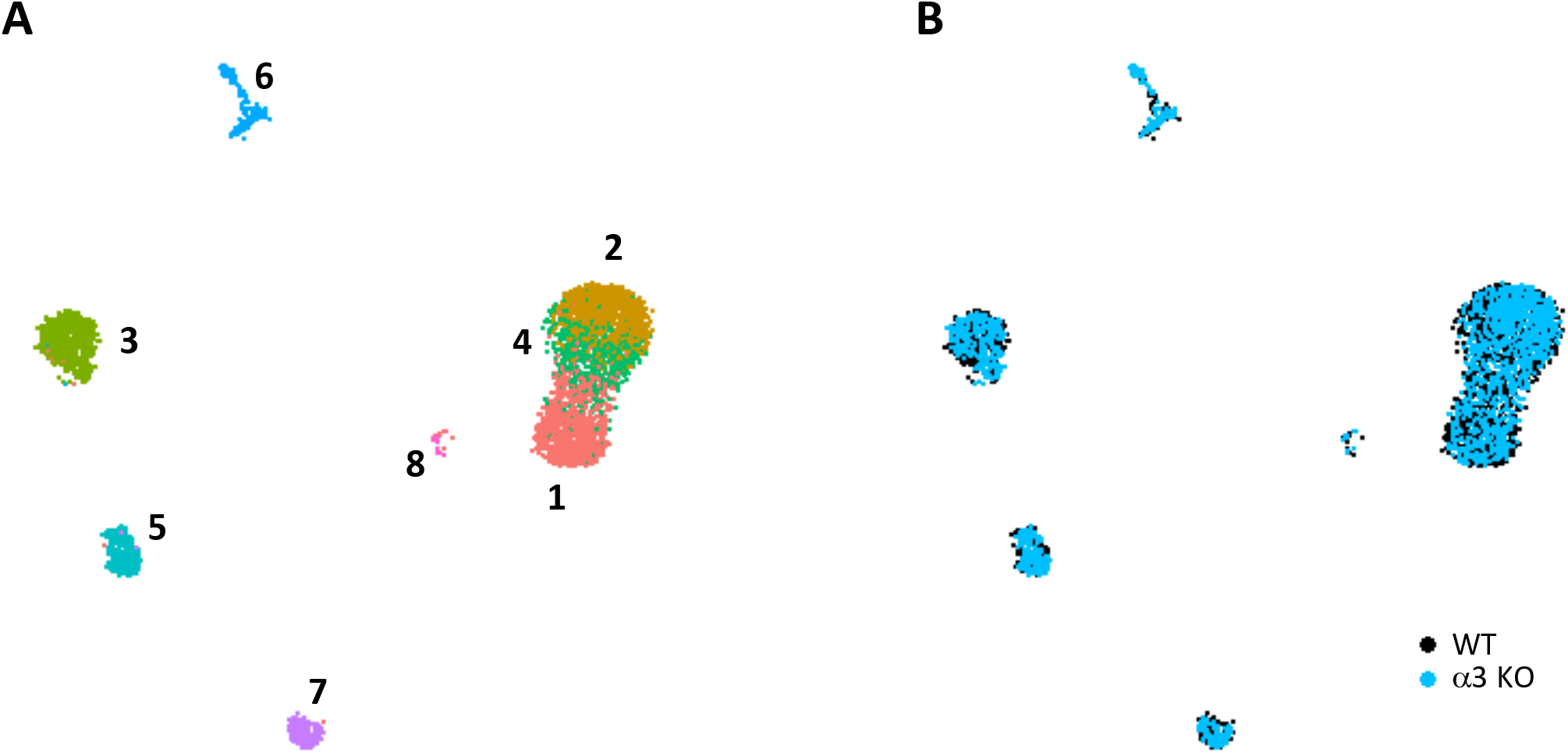
Gene expression profiles of non-neuronal cells from WT and α3 KO SCG do not differ significantly. (**A-B**) Non neuronal cells from WT (black) and α3 KO (blue) SCG were clustered into 8 subtypes and visualized with UMAPs sorted by (**A**) subtype and sorted by (**B**) genotype.

## Supplemental Information titles and legends

**Table S1. Differentially expressed genes in iris.**

Log2 fold change and corresponding p values for genes identified by RNAseq in iris tissue between α3 KO vs. WT, and between mosaic vs. WT. Genes with low counts were filtered out. Values highlighted in green are not significantly different from WT (log2FC<0.585, p value < 0.05).

**Table S2. Selected differentially expressed genes in iris.**

Log2 fold change and corresponding p values for selected genes identified by RNAseq in iris tissue between α3 KO vs. WT, and between mosaic vs. WT. Genes with low counts were filtered out. Values highlighted in green are not significantly different from WT (log2FC<0.585, p value < 0.05).

**Table S3. Marker genes for neuronal subtypes in WT SCG.**

Marker genes that define WT SCG subtypes are listed with corresponding average ln(x+1) fold change between neurons of a particular SCG subtype compared to the average expression in all other subtypes combined, p values and Bonferonni adjusted p values, the proportion of neurons expressing the given gene in that SCG subtype, and the proportion of neurons expressing the given gene in all other subtypes combined.

**Table S4. Differentially expressed genes in SCG neurons.**

Genes that are expressed at significantly different levels between WT and α3 KO neurons in at least one SCG subtype are listed. For each SCG subtype, the table contains normalized average expression levels in WT and α3 KO neurons, the average ln(x+1) fold change between WT and α3 KO neurons, and the normalized average expression level in X^RFP^ neurons. Values in bold indicate significantly different expression between WT and α3 KO in that subtype, and values in blue indicate genes that are restored towards WT levels in X^RFP^ neurons.

**Table S5. Marker genes for non-neuronal subtypes in WT SCG.**

Marker genes that define non-neuronal subtypes in WT SCG are listed with corresponding average ln(x+1) fold change between cells of a particular SCG subtype compared to the average expression in all other subtypes combined, p values and Bonferonni adjusted p values, the proportion of cells expressing the given gene in that SCG subtype, and the proportion of cells expressing the given gene in all other subtypes combined.

**Table S6. Differentially expressed genes in SCG non-neuronal cells.**

The table lists genes that are expressed at significantly different levels between non-neuronal cells in WT and α3 KO SCG, with corresponding average ln(x+1) fold change between cells of a particular non-neuronal subtype compared to the average expression in all other subtypes combined, p values and Bonferonni adjusted p values, the proportion of cells expressing the given gene in that non-neuronal subtype, and the proportion of cells expressing the given gene in all other non-neuronal subtypes combined.

